# Mindfulness meditators do not show differences in electrophysiological measures of error processing

**DOI:** 10.1101/438622

**Authors:** Neil W Bailey, Kavya Raj, Gabrielle Freedman, Bernadette M Fitzgibbon, Nigel C Rogasch, Nicholas T Van Dam, Paul B Fitzgerald

**Affiliations:** Monash Alfred Psychiatry Research Centre, Monash University Central Clinical School, Commercial Rd, Melbourne, Victoria, Australia.; Brain and Mental Health Research Hub, Monash Institute of Cognitive and Clinical Neurosciences and Monash Biomedical Imaging, Monash University, Clayton, 3168 VIC, Australia.; School of Psychological Sciences, The University of Melbourne, Parkville, VIC, Australia; Department of Psychiatry, Icahn School of Medicine at Mount Sinai, New York, NY, USA; Epworth Healthcare, The Epworth Clinic, Camberwell, Victoria, Australia, 3004.

**Keywords:** Mindfulness, Meditation, EEG, Error Processing, ERN, Pe

## Abstract

Mindfulness meditation may improve attention and self-regulation. One component of attention and self-regulation that may allow these improvements is performance monitoring. Neural correlates of performance monitoring can be objectively measured with electroencephalogram (EEG) via the error related negativity (ERN) and error positivity (Pe). Previous research assessing the ERN and Pe in meditators has resulted in inconsistent findings; some have reported alteration in peak amplitudes from both very brief meditation practice and long-term meditation practice, while others have failed to provide evidence for differences in the ERN or Pe. However, recently developed EEG analysis techniques allow for more rigorous analyses than have been used in past investigations. The current study measured the ERN and Pe, as well as post-error alpha suppression, during a Go/Nogo task, and emotional and colour Stroop tasks. The measures were compared between 22 experienced meditators (mean of 8 years of practice) and 20 healthy controls. The results suggested no differences in the ERN, Pe, or post-error alpha suppression (all p > 0.05), even when varying multiple analysis parameters. The study showed equivalent statistical power to previous research, and > 85% power to detect medium effect sizes present in previous research. Bayes Factor analysis indicated the null hypotheses were > 3.5 more likely than any of the alternative hypotheses for the ERN or Pe. These results suggest that meditation may not alter neural activity related to error processing, despite prior research suggesting that it does.

## Introduction

Studies suggest that mindfulness meditation may improve cognitive functions as a result of enhanced attentional capacities and self-regulatory skills (Jha, Krompinger, & Baime, 2007; Tang et al., 2007; Teper, Segal, & Inzlicht, 2013). One important component of both attentional function and self-regulation is performance monitoring, which ensures behaviour is adjusted to meet task demands or goals (Shenhav, Botvinick, & Cohen, 2013; Teper & Inzlicht, 2012; Teper et al., 2013). Optimal performance monitoring requires both awareness of behavioural performance and non-catastrophic reactions to errors, in order to allow adaptive adjustments to ensue (Bing-Canar, Pizzuto, & Compton, 2016). Mindfulness meditation may directly enhance performance monitoring, as the practice requires monitoring of the focus-of-attention and redirection back to a chosen focus when attention wanders, while maintaining an attitude of non-judgemental acceptance and/or discernment (Bishop et al., 2004; Teper & Inzlicht, 2012; Van Dam et al., 2018). Supporting this theory, the practice has been shown to alter prefrontal brain regions associated with attentional control and performance monitoring (Allen et al., 2012; Froeliger, Garland, Modlin, & McClernon, 2012; Hasenkamp & Barsalou, 2012; Tang, Hölzel, & Posner, 2015). Within the frontal regions, mindfulness meditation has been shown to increase blood flow in the anterior cingulate cortex (ACC) and the dosolateral prefrontal cortex (DLPFC) (Tang et al., 2015). Both of these regions have been have been associated with performance monitoring (Kerns et al., 2004; Shenhav et al., 2013). Error processing performed in these areas produces activity that can be measured at the scalp using electroencephalography (EEG), which is more sensitive to dynamic change than fMRI and can provide a measure of neural activity time-stamped to errors with millisecond resolution. In particular, error processing can be measured with EEG via the error related negativity (ERN) (Gehring, Goss, Coles, Meyer, & Donchin, 1993) and error positivity (Pe) (Michael Falkenstein, Hoormann, Christ, & Hohnsbein, 2000).

The ERN is a negative voltage deflection over frontal brain regions 50 to 150 ms following an error (Brázdil, Roman, Daniel, & Rektor, 2005; Michael Falkenstein et al., 2000; Van Veen & Carter, 2002). Source localization analyses suggests that ERN activity is produced by the ACC (Dehaene, Posner, & Tucker, 1994). While there are different views on the particular function of the ERN, all views suggest an underlying function for the modulation of attentional processes following an error (Friedman, 2012; Gehring, Liu, Orr, & Carp, 2012; Van Noordt, Campopiano, & Segalowitz, 2016; van Noordt, Desjardins, & Segalowitz, 2015). More specifically, the ERN has been suggested to reflect a range of performance monitoring functions including an operant conditioning signal, negative emotional response to an error, or an initial indicator of conflict between response and task demands (Bartholow, Henry, Lust, Saults, & Wood, 2012; Botvinick, Braver, Barch, Carter, & Cohen, 2001; Hoffmann & Falkenstein, 2012; Olvet & Hajcak, 2008; Van Veen & Carter, 2002; Yeung, 2004).

Larger ERN amplitudes are related to better executive and attentional function (Larson & Clayson, 2011) and improved ability to cope with stress (Compton et al., 2008).

In contrast to the ERN, the Pe is a positive voltage deflection over centro-parietal regions, occurring ~200 to 400 ms after an error, and is thought to be generated by the cingulate cortex and the insula (Herrmann, Römmler, Ehlis, Heidrich, & Fallgatter, 2004; O’connell et al., 2007; Overbeek, Nieuwenhuis, & Ridderinkhof, 2005; Ullsperger, Harsay, Wessel, & Ridderinkhof, 2010; Vocat, Pourtois, & Vuilleumier, 2008). The function of the Pe is argued to reflect conscious processing / awareness of the error, or the direction of attention towards the relevance of the error for motivational factors (Endrass, Klawohn, Preuss, & Kathmann, 2012; Hughes & Yeung, 2011; Nieuwenhuis, Ridderinkhof, Blom, Band, & Kok, 2001; Ridderinkhof, Ramautar, & Wijnen, 2009; Shalgi, Barkan, & Deouell, 2009). Increased Pe amplitudes are related to increased affective response to the error (Davies, Segalowitz, Dywan, & Pailing, 2001; M Falkenstein, 2004; Overbeek et al., 2005). Therefore, while the ERN may reflect more early attentional-based processes, the Pe may reflect the later conscious or affective response to the error.

Lastly, alpha activity seems to reflect the disengagement of brain regions, so the suppression of alpha activity following errors has been suggested to indicate increased arousal or attention in order to maintain task performance (Carp & Compton, 2009; Compton, Bissey, & Worby-Selim, 2014; Compton, Hofheimer, & Kazinka, 2013; Navarro-Cebrian, Knight, & Kayser, 2013; van Driel, Ridderinkhof, & Cohen, 2012). Post-error alpha suppression is associated with increased post-error slowing, although not increased performance (Carp & Compton, 2009).

Given the role of the ERN and Pe in attention, and their potential neural basis in the ACC and anterior insula (neural areas known to be affected by meditation) (Fox et al., 2016), these neural markers of error processing are potentially important indicators of change in relation to meditation. Similarly, post-error alpha suppression might be a marker improved ability in meditators to modulate neural activity to improve performance following an error. While a variety of changes to the ERN and Pe have been observed in relation to meditation, the findings are inconsistent (see Table 1). For example, a university student cohort exposed to a brief single session of mindful breathing training showed a reduced Pe amplitude but no changes to the ERN (Larson, Steffen, & Primosch, 2013). In contrast, brief single session training in mindful awareness of emotions lead to increased differences in the ERN window between error and correct trials (but no changes in the Pe window) (Saunders, Rodrigo, & Inzlicht, 2016). Within this same study, training to mindful awareness of thoughts showed no changes to either the ERN or Pe (Saunders et al., 2016). Bing-Canar et al. (2016) showed no differences in ERN and Pe amplitudes in university students exposed to a brief single session mindful breathing training. They did however find that their mindful breathing group showed an increase in post-error alpha suppression from baseline to post-training (Bing-Canar et al., 2016).

**Table 1.**
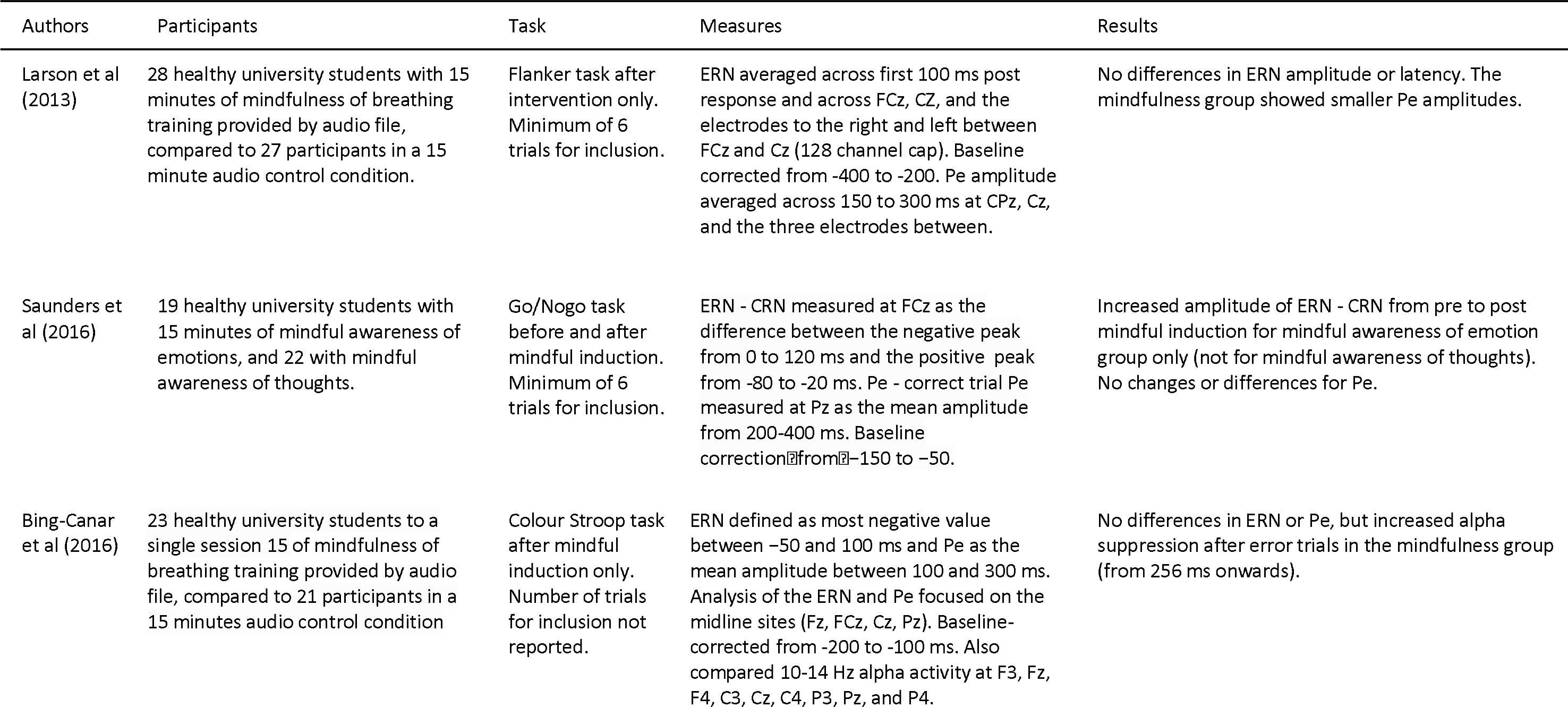
Previous research examining error processing using EEG in mindfulness meditation

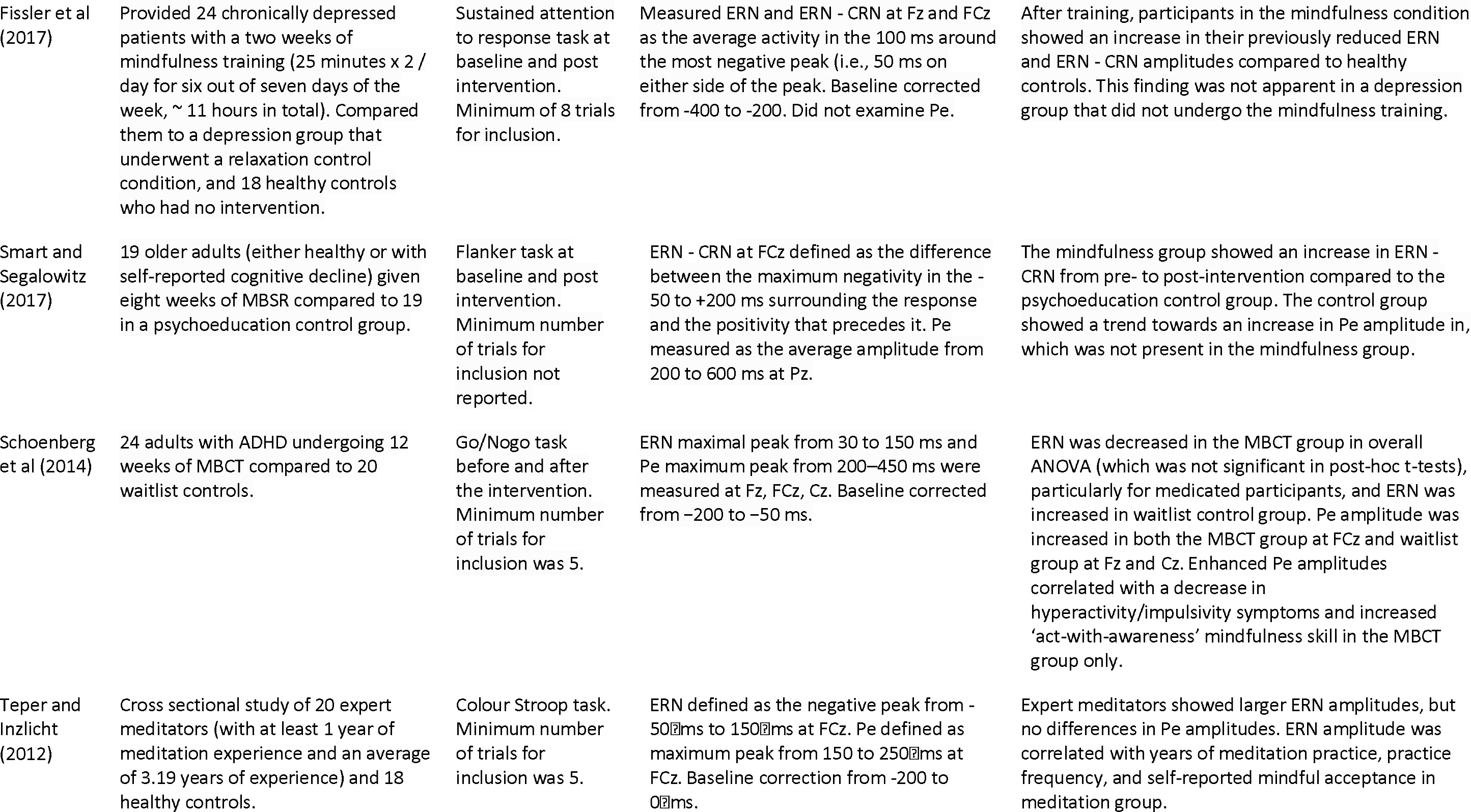

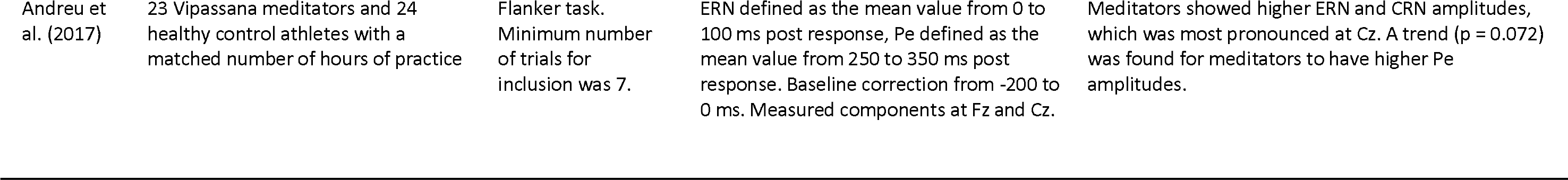

Clinical or aging populations undergoing more prolonged meditation training have shown similarly inconsistent results. Fissler et al. (2017) found that patients with depression who underwent two weeks of meditation training showed an increase to their previously reduced difference in error minus correct ERN amplitudes. These results were not found in the comparison group, who did not receive the intervention. Similarly, after an eight-week mindfulness-based stress reduction (MBSR) program, older adult participants showed an increase in the difference between error and correct ERN amplitudes compared to a psychoeducation control group (Smart & Segalowitz, 2017). However, twelve weeks of mindfulness based cognitive therapy (MBCT) for adults with ADHD showed reduced difference between error and correct ERNs, compared to an increase in the waitlist control group (Schoenberg et al., 2014). The participants in the meditation group also showed increased Pe amplitudes, which were related to increased self-reported awareness and decreased hyperactivity and impulsiveness.

Only two studies have examined error processing in experienced meditators with significant meditation experience using a cross-sectional study design. Firstly, Teper and Inzlicht (2012) studied participants who had on average 3.19 years of practice. Compared to non-meditators, the experienced meditators showed increased ERN amplitudes, which related to increased self-reported emotional acceptance. No differences were present in Pe amplitudes. Both years of meditation experience and frequency of practice correlated with ERN amplitudes, suggesting more meditation practice was related to larger ERN amplitude. Similarly, Andreu et al. (2017) found larger ERN amplitudes across both correct and incorrect trials in Vipassana meditators (average 2500 hours of meditation practice), compared to a control group of athletes with an amount of exercise comparable to the hours of meditation in the other group. There were no differences in Pe amplitude. There were also no correlations between experience and ERN.

Methodological issues may have contributed to the inconsistent findings of the research to date. In particular, all prior studies focus on single electrodes and time windows of interest. Constraining analyses with a priori assumptions in this way may miss effects of interest. In particular, single electrode analyses cannot differentiate altered strength of neural response and altered distributions of neural activity (Koenig, Kottlow, Stein, & Melie-García, 2011). Fissler et al. (2017) suggested that altered distributions of activity were present in their meditation group after visual inspection of their data, suggesting the need for analyses that examine differences in the distribution of neural activity. Additionally, subjective selections of single electrodes and time windows for analyses may inflate false positive rates (Kilner, 2013). Analysis techniques are now available that include all electrodes and time windows in the analysis, while accurately controlling the false positive rate (Koenig et al., 2011). These approaches have the advantage of not requiring a priori hypotheses regarding either where or when effects are expected and are therefore appropriate for exploratory analyses. Additionally, the majority of previous research has focused on relatively inexperienced mindfulness meditators (in some cases, as few as 15 minutes of practice; Larson et al. 2013, Saunders et al. 2016), usually no more than twelve weeks of practice (Smart and Segalowitz, 2017, Schoenberg et al. 2014). As such, it is not clear what effects prolonged practice may have on neural activity related to error processing.

Therefore, the aim of the current research was to determine whether a healthy participant sample with significant meditation experience showed alterations to the ERN and Pe, as well as to alpha suppression following errors. We hypothesised that meditators would show increased ERN amplitudes, in line with the findings of Teper and Inzlicht (2012) and Andreu et al. (2017). Although the Pe is thought to reflect awareness of errors, a process that might be considered to be enhanced by mindfulness meditation, we did not hypothesize a particular direction of effect for this component, as the majority of research has shown no differences, and the two studies that have shown differences showed those differences in opposite directions. Lastly, we hypothesized an increase in error related alpha suppression in the meditation group, in line with the results of Bing-Canar (2016).

## Methods

### Participants

Thirty-four meditators and 36 healthy control non-meditators were recruited via community and meditation centre advertising. The meditation group was required to have practiced meditation for a minimum of six months (a mean of 8.72 years, and 5.74 hours per week of current practice). Potential participants were screened and interviewed by trained mindfulness researchers (GF, KR, NWB), ensuring that meditation practices were consistent with Kabat-Zinn’s definition - “paying attention in a particular way: on purpose, in the present moment, and nonjudgementally” (Kabat-Zinn, 1994). Screening uncertainties were resolved by consensus between two researchers including the principal researcher (NWB). We required that meditation techniques be either focused attention on the breath or body-scan. Although the meditators included in the study came from a variety of backgrounds, requiring practices to meet these definitions ensured there would be overlap in practices which would result in similar effects and outcomes (Tang et al., 2010; Teper & Inzlicht, 2012). Participants in the control group were only included if they had no experience with any kind of meditation.

Exclusion criteria involved current or historical mental or neurological illness, or current psychoactive medication or recreational drug use. Participants were additionally interviewed with the MINI International Neuropsychiatric Interview for DSM-IV (Hergueta, Baker, & Dunbar, 1998) and excluded if they met criteria for DSM-IV psychiatric illness. Participants who scored in the mild or above range for anxiety or depression in the Beck Anxiety Inventory (BAI) (Steer & Beck, 1997) or Beck Depression Inventory II (BDI-II) (Beck, Steer, & Brown, 1996) were also excluded. All participants had normal or corrected to normal vision and were between 19 and 62 years of age.

Prior to completing the EEG task, participants provided their age, gender, years of education, handedness, and estimate of number of years, as well as number of minutes per week meditating. Participants also completed the Freiburg Mindfulness Inventory (FMI) (Walach, Buchheld, Buttenmüller, Kleinknecht, & Schmidt, 2006), Five Facet Mindfulness Questionnaire (FFMQ) (Baer, Smith, Hopkins, Krietemeyer, & Toney, 2006), BAI, and BDI-II. Table 2 summarizes these measures. All participants provided informed consent prior to participation in the study. The study was approved by the Ethics Committee of the Alfred Hospital and Monash University.

**Table 2.**
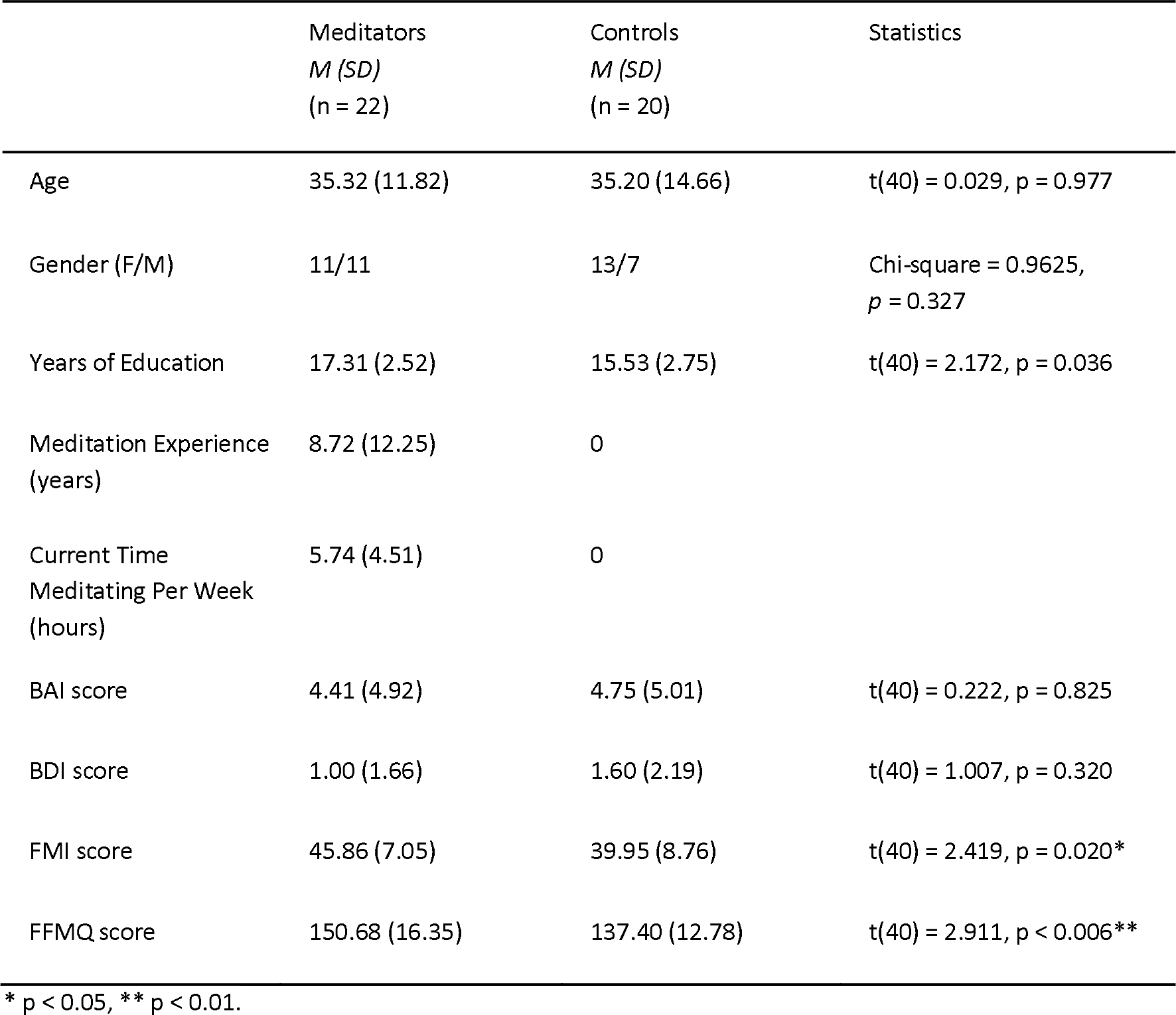
Demographic, self-report, and behavioural data.

Four controls were excluded from the study after scoring in the mild depression range on the BDI, two controls were excluding due to misunderstanding task instructions, and one control was excluded due to task non-completion. Three additional controls and three meditators were excluded from analysis due to an equipment fault. Lastly, six controls and nine meditators were excluded from analysis after providing too few accepted error related epochs for analysis (< 10). For final inclusion in analyses, 22 meditators and 20 healthy controls were included in the study, providing more than 10 errors across the Go/Nogo, colour Stroop and emotional Stroop tasks.

Although our final included participant sample size was similar to previous research, we calculated a post-hoc power analysis to determine if any null results might be explained by lack of power. Power analysis was conducted via extraction of ERN means and SDs from Teper and Inzlicht (2012); we assumed that their community recruited meditator sample would provide the most similar effects to those in our current sample (Andreu et al. 2017 recruited only Vipassana meditators). The values were input into GPower to compute post-hoc power in an independent samples t-test with a one-way tail (given our hypothesis that meditators would show a larger ERN) and alpha of 0.05 based on our sample size. We also computed power using a repeated measures ANOVA design (as used in the current study, including both error and correct trials), after converting Teper and Inzlicht (2012)’s d statistic (0.58) to f (0.29), and using the correlation between correct and incorrect ERN amplitudes from the current study (0.345).

### Procedure

Participants performed a Go/Nogo task, a colour Stroop and an emotional Stroop task (described in Bailey et al. in preparation, Raj et al. in preparation). 64-channel EEG was recorded while participants performed these tasks. A Neuroscan 64-channel Ag/AgCl Quick-Cap was used to acquire EEG through NeuroScan Acquire software and a SynAmps 2 amplifier (Compumedics, Melbourne, Australia). Electrodes were referenced to an electrode between Cz and CPz. Eye movements were recorded with vertical and horizontal EOG electrodes. Electrode impedances were kept below 5kΩ. The EEG was recorded at 1000Hz, with an online bandpass filter of 0.1 to 100Hz. Data were analysed offline in MATLAB (The Mathworks, Natick, MA, 2016a) using EEGLAB for pre-processing (sccn.ucsd. edu/eeglab) (Delorme & Makeig, 2004). Second order Butterworth filtering was applied to the data with a bandpass from 1–80 Hz and also a band stop filter between 47–53 Hz. Error and correct response trials were re-coded, and data were epoched from −500 to 1500 ms surrounding the onset of the stimulus presentation for each trial. Epochs were visually inspected by an experimenter experienced with EEG analysis and blinded to the group of each participant, and periods containing muscle artefact or excessive noise were excluded, as were channels showing poor signal.

Data were combined with epoched data from correct responses from each task (results of which are being prepared in a separate publication) for Independent Component Analysis (ICA) processing, to provide ICA with more data for accurate component selection. Adaptive Mixture ICA (AMICA) (Palmer, Makeig, Kreutz-Delgado, & Rao, 2008) was used to manually select and remove eye movements and remaining muscle activity artefacts. After artifactual ICA components were rejected, raw data were re-filtered from 0.1-80 Hz, all previous channel and epoch rejections were applied, and rejected ICA components were applied to this 0.1-80 Hz filtered data to avoid rejecting low frequency brain activity around 1 Hz (prior to ICA rejection, data below 1 Hz was filtered out as it adversely impacts the ICA process). Rejected electrodes were re-constructed using spherical interpolation (Perrin, Pernier, Bertrand, & Echallier, 1989). Data were then visually inspected again to ensure the artefact rejection process was successful. Recordings were re-referenced offline to an averaged reference and baseline corrected to the −400 to −100 ms period, and epochs from each condition and participant were averaged for ERP analyses.

### Measures

Previous literature shows a lack of consensus as to the most reliable number of accepted error epochs required for analysis of the ERN and Pe. While some researchers have recommended 6 epochs as a minimum for valid and reliable analysis, other researchers have suggested minimums of 14-15 (Fischer, Klein, & Ullsperger, 2017; Larson, Baldwin, Good, & Fair, 2010; Olvet & Hajcak, 2009; Rietdijk, Franken, & Thurik, 2014; Steele et al., 2016). As such, we chose to include participants only if they provided a minimum of 10 accepted error epochs, in order to err on the side of caution with regards to validity and reliability (as 10 epochs has been found to be the threshold for high internal reliability (Olvet & Hajcak, 2009)), while still including enough participants to obtain sufficient power to detect potential significant differences. In addition to the error response related epochs, we also extracted a matched number of correct response related epochs from each task. Each accepted error response related epoch had a correct response related epoch selected from the same participant and task. For the Go/Nogo task, error responses were responses made to Nogo trials, and correct responses were made to Go trials. For the Stroop tasks, error responses were button presses other than with the button paired with the stimulus being presented (for example, a press of button 2, when instructions stated the stimulus being presented was paired with button 1). Additionally, the correct response related epoch with the smallest difference in reaction time to the error response reaction time was selected to ensure correct and error response related epochs were matched for reaction time. We also attempted to determine if the meditation group showed higher alpha suppression in error trials compared to correct trials than the control group, which Bing-Canar et al. (2016) suggested to be related to increased attentional engagement following an error in order to ensure correct response in the following trials. Data from all electrodes in each epoch was subjected to a fast Fourier transform (FFT) using a cosine windowing method from 10-14 Hz (the alpha range used by Bing-Canar et al. 2016).

### Data Analysis

#### replication comparisons

In addition to the whole scalp analysis with the Randomisation Graphical User Interface (RAGU), more traditional single electrode comparisons were planned for comparison with previous research. Data from midline electrodes Fz, FCz, Cz, CPz, and Pz had activity averaged during the ERN window (defined as activity from 50 to 150 ms following the response) and the Pe window (defined as activity from 200 to 400 ms following the response). These averaged windows were calculated for both correct and error responses. SPSS version 23 was used for frequentist analyses of single electrode data, and Bayesian analyses were performed using JASP. Repeated measures ANOVAs were used to conduct a 2 group (meditators vs controls) × 2 trial type (correct vs error) × 5 electrode (Fz, FCz, Cz, CPz, Pz) comparison for the ERN and Pe separately. Similarly, separate repeated measures ANOVAs were conducted for alpha suppression values averaged across two time periods of interest, in direct replication of the comparisons of Bing-Canar et al. (2015) (0 to 256 ms and 256 to 512 ms). These comparisons involved a 2 group (meditators vs controls) × 2 trial type (correct vs error) × 9 electrode (F3, Fz, F4, C3, Cz, C4, P3, Pz, P4). All sphericity violations were addressed via the Greenhouse-Geisser correction (Greenhouse & Geisser, 1959). Bayes Factor repeated measures ANOVA analyses were used to determine the likelihood of the null hypothesis in contrast to the alternative hypothesis where null results were found in frequentist statistics (Rouder, Morey, Verhagen, Swagman, & Wagenmakers, 2017). The suggested comparison between models containing a hypothesized effect to equivalent models stripped of the effect (excluding higher order interactions) was performed for these analyses.

Lastly, repeated measures ANOVAs were used to compare the number of accepted epochs from each task from each group. Repeated measures ANOVAs were used to compare reaction times between groups and across correct/incorrect trials for each task separately (because not all participants contributed errors from each task). The same repeated measures ANOVAs were conducted for post-error and post-correct reaction times in order to examine whether post-error slowing was different between groups. Independent samples t-tests were used to compare the total number of errors from each group, demographic variables, and self-report variables between groups (except for gender ratios in each group, which was compared with a Chi-squared test). Lastly, because neural and behaviour data showed null results, an exploratory repeated measures ANOVA was used to compare groups in the two FMI measures that seem most related to subjective experience in response to errors, in an attempt to assess whether subjective experience of errors might differ between the groups. The items were “I see my mistakes and difficulties without judging them” and “I am friendly to myself when things go wrong”.

#### primary comparisons

Due to the limitations of selected electrode and timepoint analyses, primary EEG data statistical comparisons were conducted using the RAGU to compare scalp field differences across all electrodes and time points with permutation statistics without making any a priori assumptions about electrodes or windows for analysis (Koenig et al., 2011). This reference-free method takes advantage of the additive nature of scalp fields to allow comparisons of neural activity between groups and conditions without estimation of active sources by calculating a difference scalp field. This difference scalp field shows the scalp field of brain sources that differed between the two groups / conditions, while brain sources that did not differ result in zero scalp field difference (Koenig et al., 2011). RAGU controls for multiple comparisons in both time and space using permutation statistics (see Koenig et al. 2011 and supplementary materials for details).

RAGU also allows for independent comparisons of overall neural response strength (with the global field power - GFP test) and distribution of neural activity (with the Topographic Analysis of Variance - TANOVA). Prior to the TANOVA, a Topographical Consistency Test (TCT) was conducted to ensure a consistent distribution of scalp activity within each group / condition.

GFP and TANOVA tests were used to conduct repeated measures ANOVA design statistics, examining 2 group (meditators vs controls) × 2 condition (corrects vs errors) comparisons for data from −100 to 700 ms surrounding the response. Five thousand permutations were conducted with an alpha of p = 0.05. To compare alpha suppression between groups (as per Bing-Canar et al. 2016), we used a repeated measures ANOVA design in RAGU to conduct a 2 groups (meditators vs controls) × 2 condition (corrects vs errors) with Root Mean Square (RMS) and TANOVA tests (to separately compare overall neural response strength and distribution of neural activity respectively). It should be noted that when frequency comparisons are computed with RAGU, the average reference is not computed on the frequency transformed data (the average reference was computed prior to the frequency transforms). As such, the test is a comparison of the RMS between groups, a measure which is a valid indicator of neural response strength in the frequency domain. In other respects, the statistic used to compare RMS between groups is identical to the GFP test described in the previous section (T. Koenig 2018, Department of Psychiatric Neurophysiology, University Hospital of Psychiatry, personal communication). Post-hoc GFP and TANOVA tests averaged across time periods of significance after global duration multiple comparison controls were planned to explore any significant effects.

## Results

### Demographic and Behavioural Data

Demographic and self-report differences are presented for the participants included in the neural analysis, as the neural analysis was the main focus of the study. Results are summarised in table 2. For participants included in the neural analysis, no significant differences were present between groups in age, BAI, BDI, gender or handedness (all p > 0.3). Meditators showed significantly more years of education [t(40) = 2.172, p = 0.036], higher FMI [t(40) = 2.419, p = 0.020], and FFMQ scores [t(40) = 2.911, p < 0.006]. All participants in the meditation group except one had more than two years of experience. Meditators also showed significant differences to controls in the two FMI measures related to subjective experience following the commission of errors: “I see my mistakes and difficulties without judging them” (meditator mean = 3.227 SD = 0.528, control mean = 2.842, SD = 0.834) and “I am friendly to myself when things go wrong” (meditator mean = 3.182 SD = 0.588, control mean = 2.526, SD = 0.697) [F(1,40) = 9.020, p = 0.005, partial eta squared = 0.188].

Percentage correct and reaction times across all trials are reported in other studies (Bailey et al. in press, Raj et al. in press). No main effect of group or interaction between group (meditators vs controls) and task (Go/Nogo, colour Stroop, emotional Stroop) was found in comparisons of the number of accepted error epochs, nor were there differences in the total number of error epochs included between groups (all p > 0.05).

Results of RT comparisons showed no significant differences between groups or interaction between group and correct/incorrect responses (all p > 0.10), with the exception of an interaction between group and correct/incorrect responses for the emotional Stroop task, where controls showed a longer RT for correct than incorrect responses, while meditators showed no differences [F(1,36) = 4.168, p = 0.049, partial eta squared = 0.104]. However, Box’s test showed non-equality of covariance matrices for this comparison (p = 0.008), which may have inflated the significance. Post-hoc t-tests indicated this interaction was not due to between group differences in either trial type (both p > 0.2), and could be accounted for by a larger difference between correct and error reaction times in controls [t(17) = 1.791, p = 0.091, a 13.76 ms difference], but not in meditators [t(19) = −0.820, p = 0.422]. Results can be viewed in table 3. Lastly, there was an main effect of post error slowing, with both groups showing slower RTs after errors [F(1,40) = 21.649, p < 0.001]. However, no interaction was found between group and RT following correct vs error trials for any task, suggesting meditators did not show more post-error slowing than controls (all p > 0.10).

**Table 3.** Behavioural and accepted epoch data.

|  | Meditators<br>(n=22) | Controls<br>(n=20) | t Statistic | F Statistic |
| --- | --- | --- | --- | --- |
| Number of Accepted Nogo Error Epochs | 7.64 (4.67) | 7.20 (5.79) | $t(37) = 0.270, p = 0.789$ | Group x task interaction:<br>$F(1,40) = 0.512, p = 0.601$ |
| Number of Accepted Colour Stroop Error Epochs | 4.27 (2.39) | 5.25 (7.96) | $t(36) = 0.550, p = 0.586$ | |
| Number of Accepted Emotional Stroop Error Epochs | 5.09 (3.74) | 7.05 (5.81) | $t(36) = 1.312, p = 0.197$ | |
| Total Accepted Error Epochs | 17.00 (6.52) | 19.50 (10.39) | $t(40) = 0.943, p = 0.351$ | $t(40) = 0.943, p = 0.351$ |
| RT of Nogo Errors | 312.36 (52.29) | 296.66 (39.80) | $t(37) = 1.028, p = 0.311$ | Group x trial interactions:<br>$F(1,37) = 1.028, p = 0.317$ |
| RT of Go Corrects | 319.59 (54.55) | 308.19 (43.10) | $t(37) = 0.707, p = 0.484$ | |
| RT of Colour Stroop Errors | 732.60 (136.37) | 674.02 (150.16) | $t(36) = 1.258, p = 0.216$ | $F(1,36) = 0.014, p = 0.906$ |
| RT of Colour Stroop Corrects | 733.13 (136.67) | 675.28 (155.19) | $t(36) = 1.221, p = 0.230$ | |
| RT of Emotional Stroop Errors | 455.84 (93.08) | 477.58 (110.25) | $t(36) = 0.659, p = 0.514$ | $F(1,36) = 4.168, p = 0.049$ |
| RT of Emotional Stroop Corrects | 453.15 (86.31) | 491.34 (99.00) | $t(36) = 1.270, p = 0.212$ | |
| Post-error RT for Go Trials | 433.16 (81.87) | 417.32 (92.13) | $t(37) = 0.482, p = 0.634$ | $F(1,37) = 1.191, p = 0.283$ |
| Post-correct RT for Go Trials | 416.01 (62.29) | 391.28 (52.36) | $t(37) = 1.127, p = 0.270$ | |
| Post-error RT for Colour Stroop | 727.30 (146.79) | 751.62 (174.77) | $t(36) = 0.460, p = 0.648$ | $F(1,37) = 0.087, p = 0.770$ |
| Post-correct RT for Colour Stroop | 624.61 (101.34) | 635.58 (88.69) | $t(36) = 0.347, p = 0.730$ | |
| Post-error RT for Emotional Stroop | 587.62 (148.74) | 653.01 (137.17) | $t(36) = 1.317, p = 0.197$ | $F(1,36) = 0.044, p = 0.836$ |
| Post-correct RT for Emotional Stroop | 502.37 (66.56) | 558.54 (80.47) | $t(36) = 2.228, p = 0.033$ | |

### Power Analysis Showed Sufficient Statistical Power

The t-test design power analysis suggested power of 0.863, sufficient to detect significant effect sizes if they were of a similar size to Teper and Inzlicht (2012). The repeated measures ANOVA design power analysis indicated the current study had even more power (0.893). The current results showed d = 0.194 for ERN comparisons at FCz, a much smaller effect size than Teper and Inzlicht’s (2012) cohen’s d = 0.58 for the same comparison, with a similar sample size (20 meditators, 18 controls).

### Single Electrode Analyses of ERN and Pe Showed No Differences between Groups

There were no significant differences for the ERN group comparison, nor interactions between group and correct/error, nor interactions between group, correct/error, and electrode (all p > 0.35, statistics reported in table 3). Bayes Factor analysis of the ERN data indicated that the null hypothesis of no difference between groups was 5.56 times more likely to be true than the alternative hypothesis, 6.67 times more likely to be true for the interaction between group and correct/error, and 10.42 times more likely for the interaction between group, correct/error and electrode. No significant differences were found for the Pe group comparison, nor interactions involving group and response, nor interactions between group, response, and electrode (all p > 0.45, statistics reported in table 4). Bayes Factor analysis of the Pe data indicated that the null hypothesis of no difference between groups was 4.78 times more likely to be true than the alternative hypothesis, 3.83 times more likely to be true for the interaction between group and correct/error, and 23.81 times more likely for the interaction between group, correct/error and electrode. Single electrode ERP plots from these electrodes can be viewed in figure 7, and averaged activity across the ERN and Pe windows at each electrode can be viewed in figure 8 and 9 respectively. Values for each electrode, condition, and group can be viewed in table 5, waveforms can be viewed in Figure 1, and means are plotted in Figures 2 and 3.

**Table 4.** Statistics for single electrode comparisons.

|  | Null Hypothesis Test | Bayes Factor Analysis for Null Hypothesis |
| --- | --- | --- |
| **ERN** |  |  |
| Between groups | $F(1,40) = 0.002$, $p = 0.968$, partial eta squared $< 0.001$ | 5.56 |
| Group x Correct/Incorrect | $F(1,40) = 0.072$, $p = 0.790$, partial eta squared $= 0.002$ | 6.67 |
| Group x Correct/Incorrect x Electrode | $F(1,40) = 1.086$, $p = 0.365$, partial eta squared $= 0.026$ | 10.42 |
| **Pe** |  |  |
| Between groups | $F(1,40) = 0.349$, $p = 0.558$, partial eta squared $= 0.009$ | 4.78 |
| Group x Correct/Incorrect | $F(1,40) = 0.551$, $p = 0.462$, partial eta squared $= 0.014$ | 3.83 |
| Group x Correct/Incorrect x Electrode | $F(1,40) = 0.371$, $p = 0.829$, partial eta squared $= 0.009$ | 23.81 |

**Table 5.** Averaged ERN (50-150 ms) and Pe (200-400 ms) activity from single electrodes.

|  | Meditator Corrects<br>*M (SD)* | Control Corrects<br>*M (SD)* | Meditator Errors<br>*M (SD)* | Control Errors<br>*M (SD)* |
| --- | --- | --- | --- | --- |
| **ERN** |  |  |  |  |
| Fz | 0.35 (1.85) | -0.11 (3.19) | -0.56 (2.12) | -2.02 (3.45) |
| FCz | 1.94 (1.93) | 1.36 (2.44) | -0.70 (2.86) | -1.31 (3.40) |
| Cz | 2.37 (1.66) | 2.40 (1.21) | -0.33 (2.23) | -0.04 (2.55) |
| CPz | 1.84 (1.45) | 2.09 (1.71) | -0.26 (1.73) | 0.69 (2.01) |
| Pz | 0.43 (1.53) | 0.86 (2.35) | -0.33 (1.57) | 0.94 (1.75) |
| **Pe** |  |  |  |  |
| Fz | 1.20 (1.65) | 1.05 (1.81) | 1.85 (2.33) | 0.73 (2.51) |
| FCz | 1.94 (1.93) | 1.36 (2.44) | 2.92 (2.24) | 2.18 (2.66) |
| Cz | 0.96 (1.28) | 1.22 (1.00) | 3.16 (1.89) | 3.02 (2.62) |
| CPz | -0.11 (1.34) | 0.24 (1.23) | 2.27 (1.82) | 2.43 (2.38) |
| Pz | -1.48 (1.74) | -1.22 (1.40) | 0.81 (1.92) | 0.85 (1.85) |

**Figure 1.**
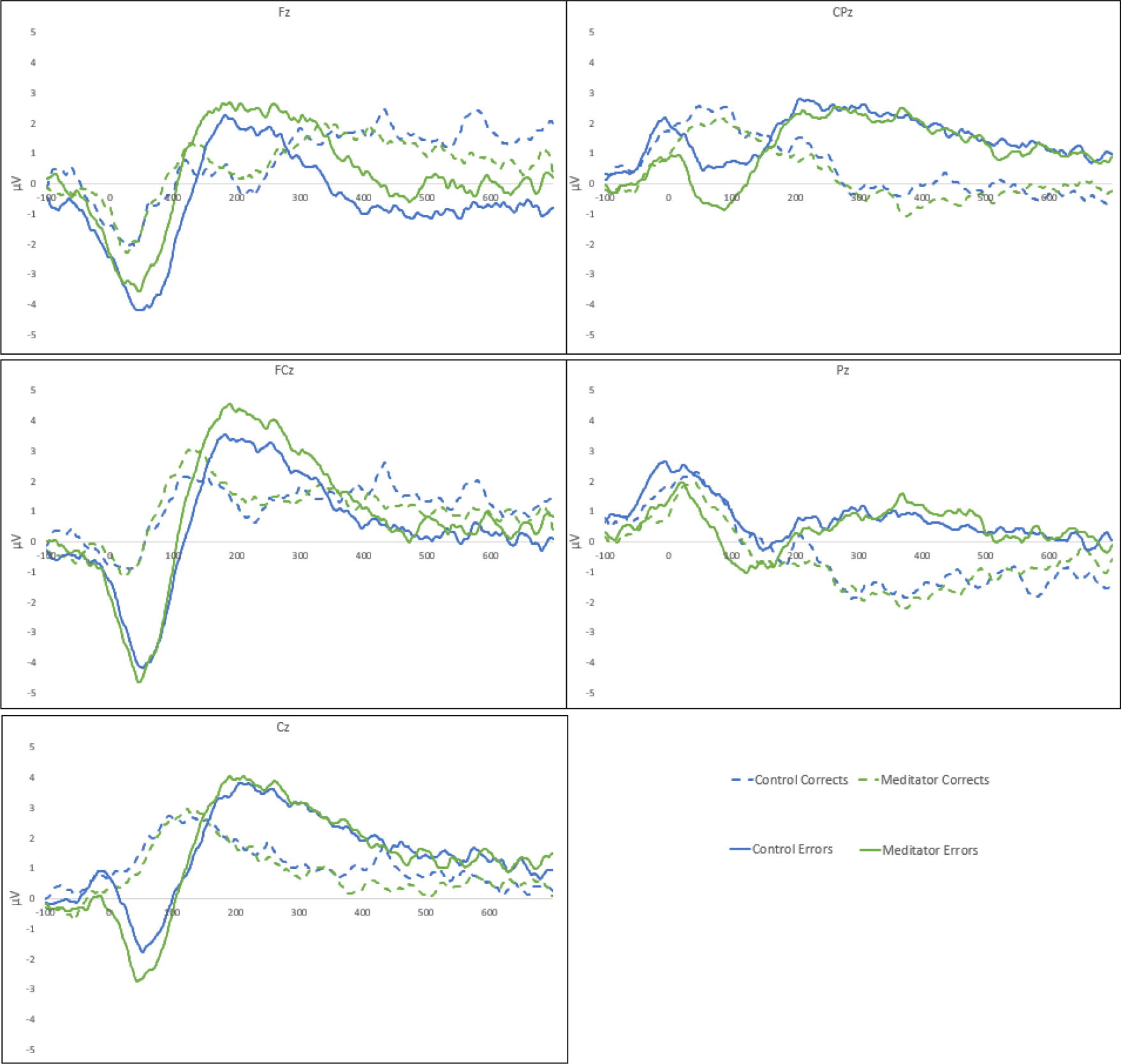
Single electrode waveforms for correct and error trials from each group.

**Figure 2.**
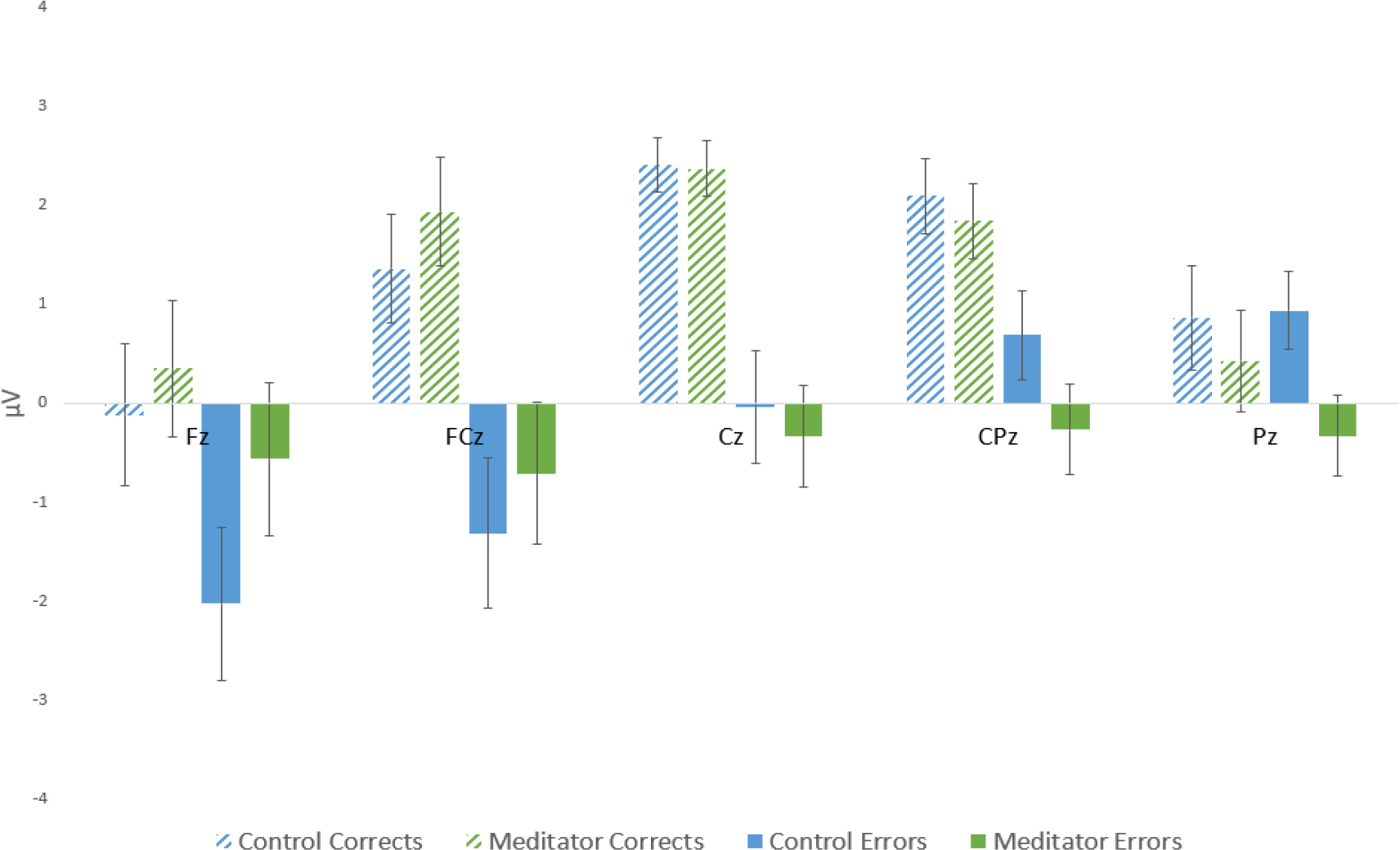
Averaged activity during the 50 to 150 ms ERN window for error and correct trials for each group. Error bars reflect standard errors. No significant differences or interactions involving group were found (all p > 0.05).

**Figure 3.**
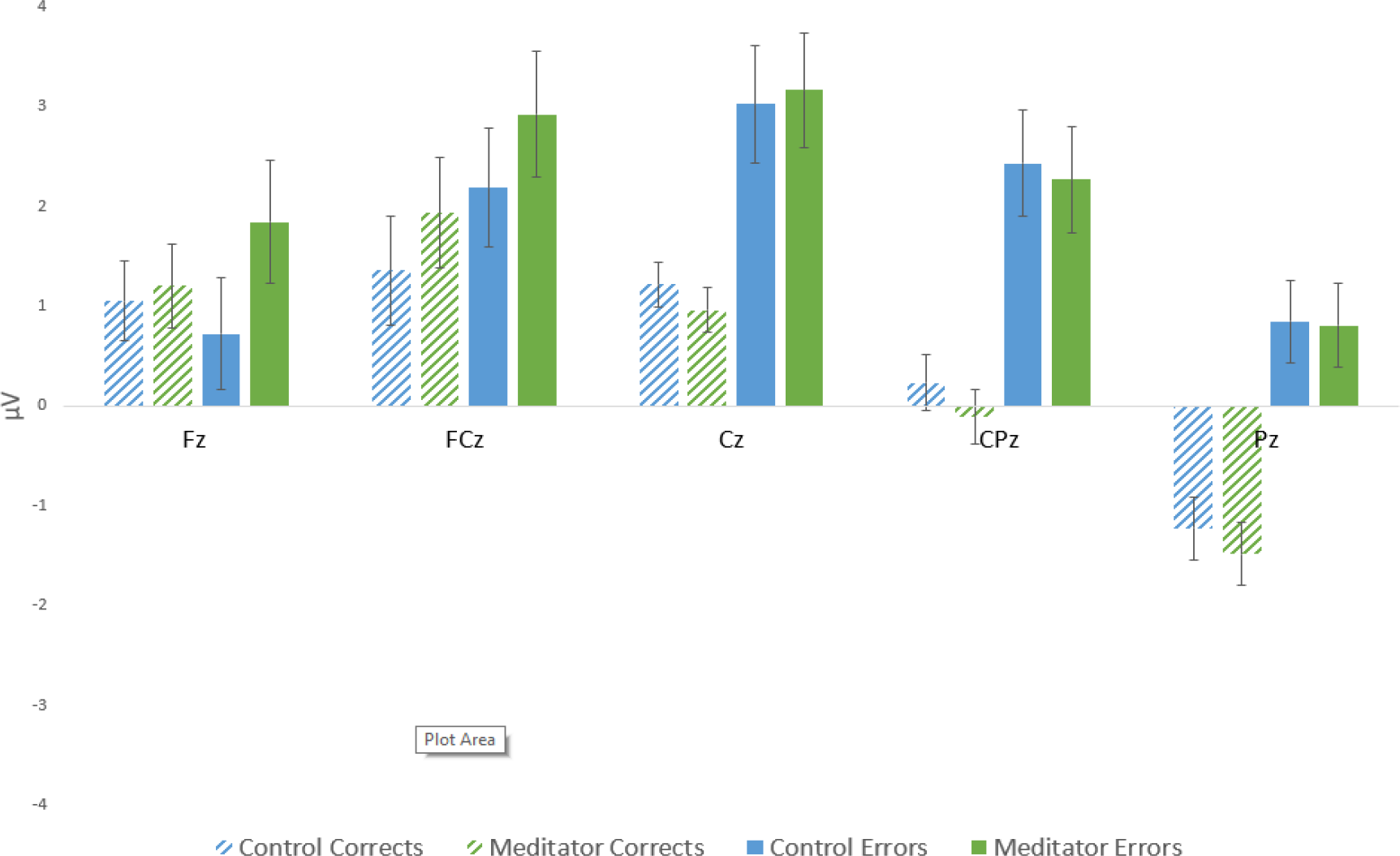
Averaged activity during the 200 to 400 ms Pe window for error and correct trials for each group. Error bars reflect standard errors. No significant differences or interactions involving group were found (all p > 0.05).

### Alpha Power Suppression in Error Trials Showed No Differences between Groups

No significant differences were found for the early alpha suppression group comparison [F(1,40) = 0.337, p = 0.565, partial eta squared = 0.008], nor interactions between group and correct/error [F(1,40) = 1.354, p = 0.252, partial eta squared = 0.033], nor interactions between group, correct/error, and electrode [F(1,40) = 0.827, p = 0.579, partial eta squared = 0.020]. No significant differences were found for the late alpha suppression group comparison [F(1,40) = 0.732, p = 0.397, partial eta squared = 0.018], nor interactions between group and correct/error [F(1,40) = 1.032, p = 0.316, partial eta squared = 0.025], nor interactions between group, correct/error, and electrode [F(1,40) = 0.194, p = 0.992, partial eta squared = 0.005]. Bayes Factor analysis of the early alpha suppression data indicated that the null hypothesis of no difference between groups was 2.27 times more likely to be true than the alternative hypothesis, and 166.67 times more likely for the interaction between group, correct/error and electrode. However, the alternative hypothesis was 1.97 times more likely to be true for the interaction between group and correct/error than the null hypothesis. Bayes Factor analysis of the late alpha suppression data indicated that the null hypothesis of no difference between groups was 2.12 times more likely to be true than the alternative hypothesis, and 62.50 times more likely for the interaction between group, correct/error and electrode. However, the alternative hypothesis was 1.28 times more likely to be true for the interaction between group and correct/error than the null hypothesis. Alpha suppression data from single trials can be viewed in Figure 4. To overcome the potential limitations intrinsic to single or selected electrode and time window analyses, we also conducted analyses including all electrodes and time points, the results of which are reported in the following section.

**Figure 4.**
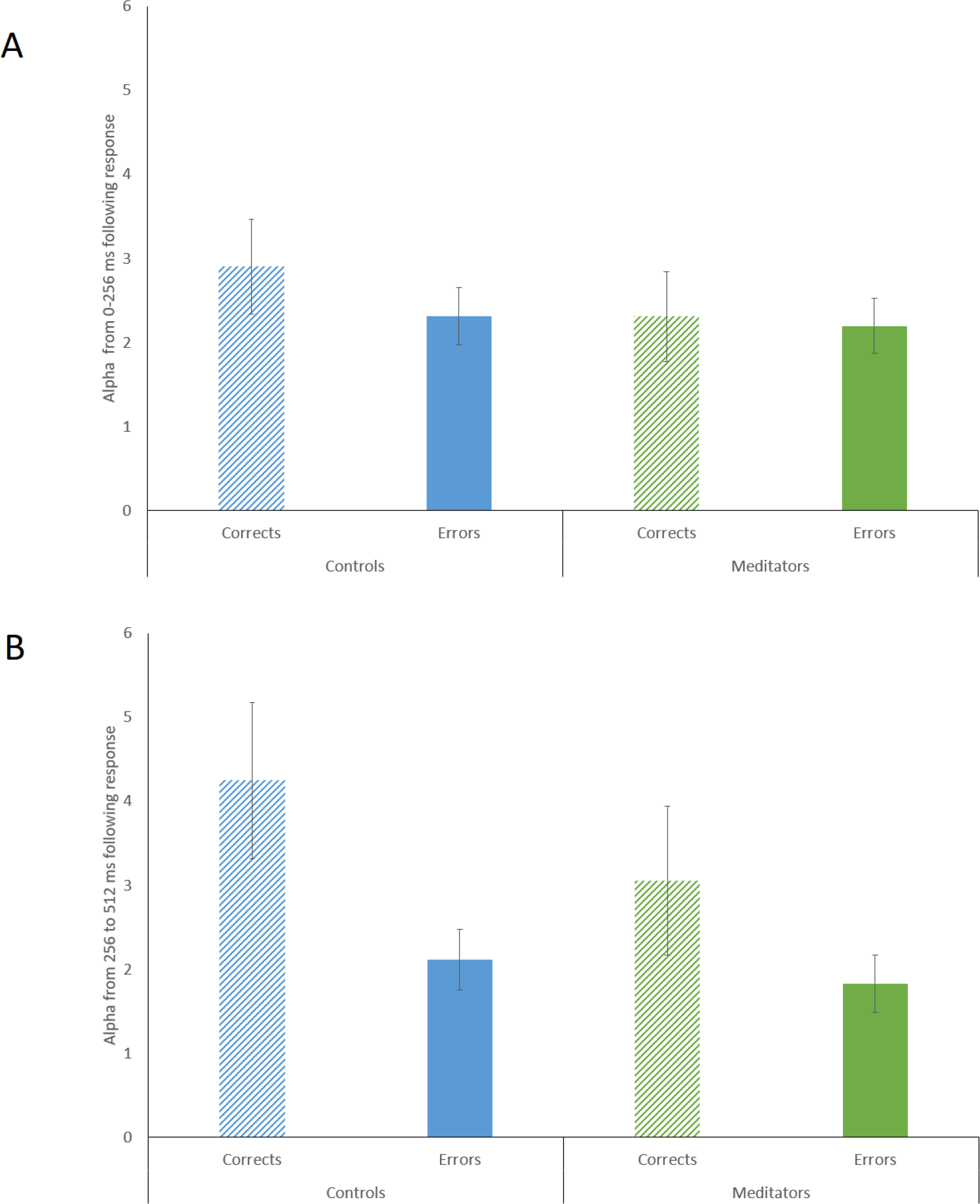
Alpha suppression values averaged across single electrodes included in the analysis (F3, Fz, F4, C3, Cz, C4, P3, Pz, P4). A – Alpha activity during the early window following the response (0-256 ms). B – Alpha activity during the later window following response (256-512 ms). Error bars reflect standard errors.

### ERN and Pe Topographical Consistency Test Showed Consistent Activity Within Groups

Both groups showed consistent topographical activation in both conditions across the epoch after responses occurred (except for a brief period around 115 ms in control error trials, see Figure 7). This indicates a consistent distribution of neural activity across the epoch within each condition and group, demonstrating that between group comparisons are valid.

### ERN and Pe Global Field Potential Test Showed No Differences Between Groups

No significant differences were shown in the GFP comparisons for the main effect of the comparison between groups nor for the group-by-correct/incorrect interaction (see Figure 5, note that p-values stayed above 0.05 for the entire epoch, except for a few short durations which did not survive the duration control for multiple comparison value of 50 ms for the group main effect and 29 ms for the interaction). However, there was a significant difference in the main effect of trial type from 172 to 293 ms (during the Pe window), where error trials showed larger GFP responses than correct trials (p = 0.001 when averaged across the window of significance, see figure 6). This period did survive duration multiple comparison control. These results indicate that neural response strength did not differ between the meditator and control group, nor that group interacted with trial type at any point across the epoch in neural response strength (including during the ERN and Pe windows). The results do however indicate that error responses showed larger neural response strength during the early Pe window across both groups.

**Figure 5.**
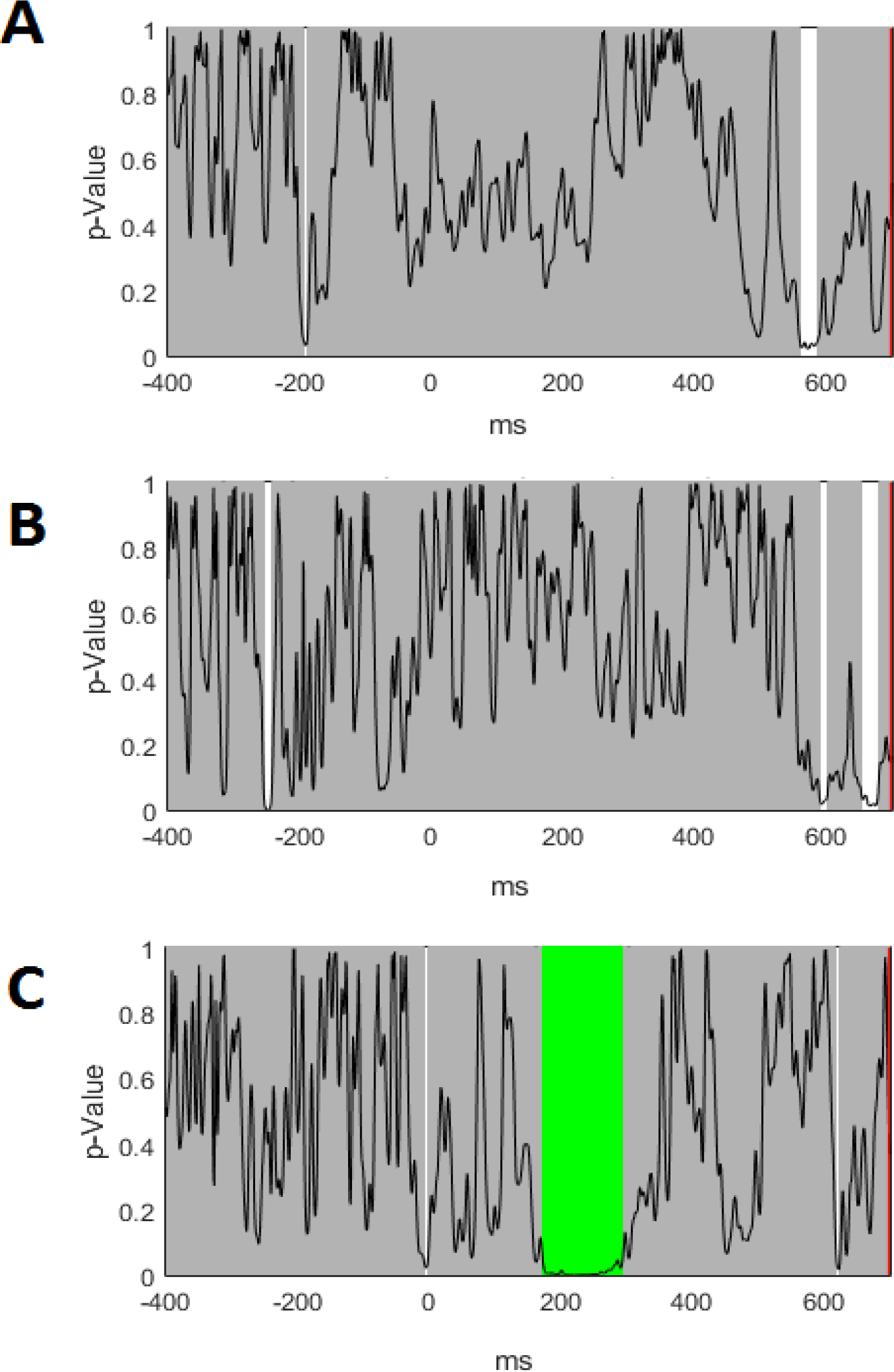
P-value graphs for GFP comparisons. The black line reflects the p-value, grey periods reflect no significant differences between groups, white periods reflect significant differences that did not survive duration multiple comparison controls, and green periods reflect significant differences that do survive duration multiple comparison controls. A - Group main effect, B - Group by correct / incorrect interaction, C - Correct / incorrect main effect (note the significant difference from 172 - 293 ms, which survived duration control for multiple comparisons).

**Figure 6.**
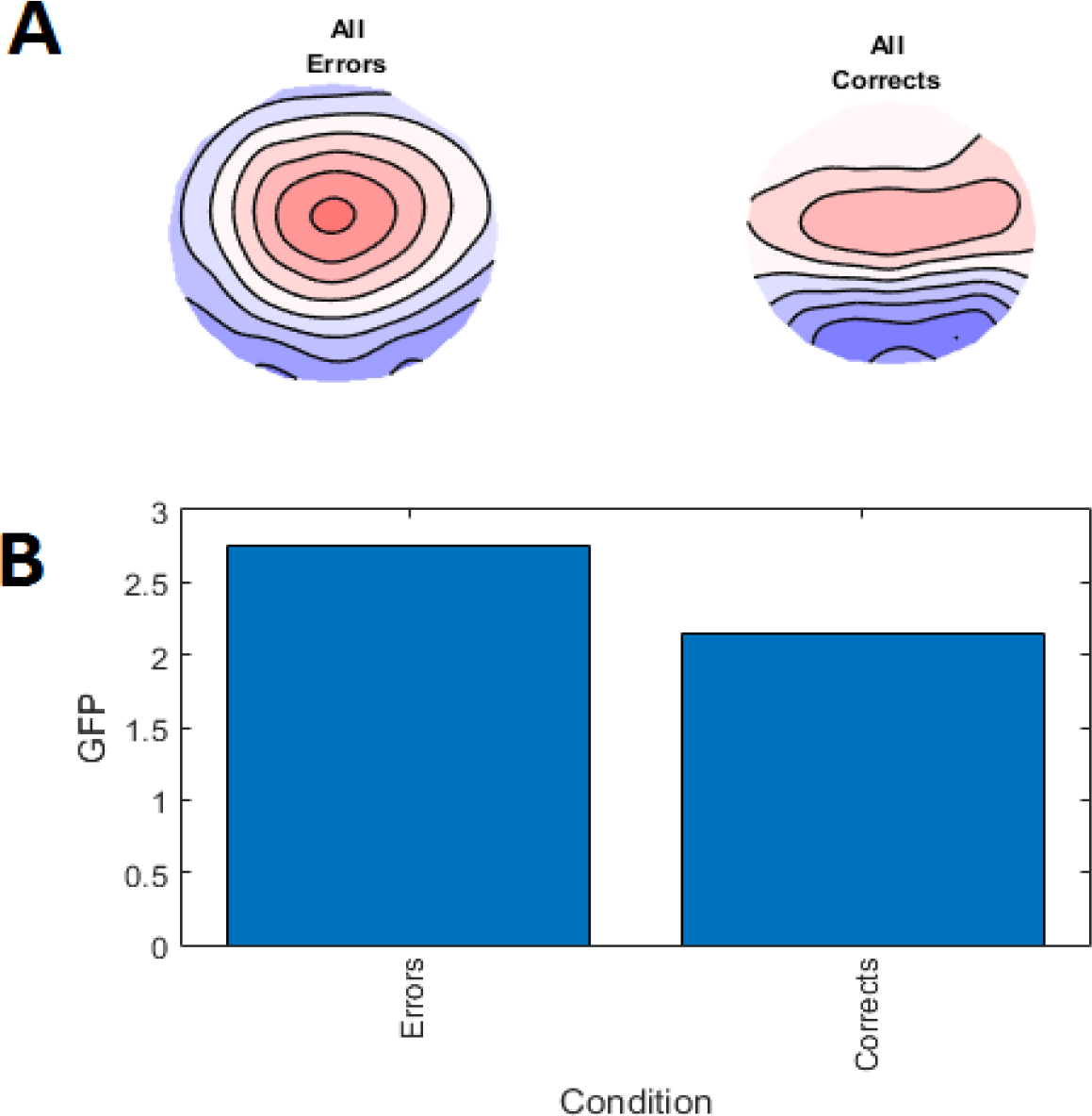
A - Averaged topographies for errors and correct trials averaged across the significant time period 172 - 293 ms. B - Averaged GFP in error and correct trials (p = 0.001).

**Figure 7.**
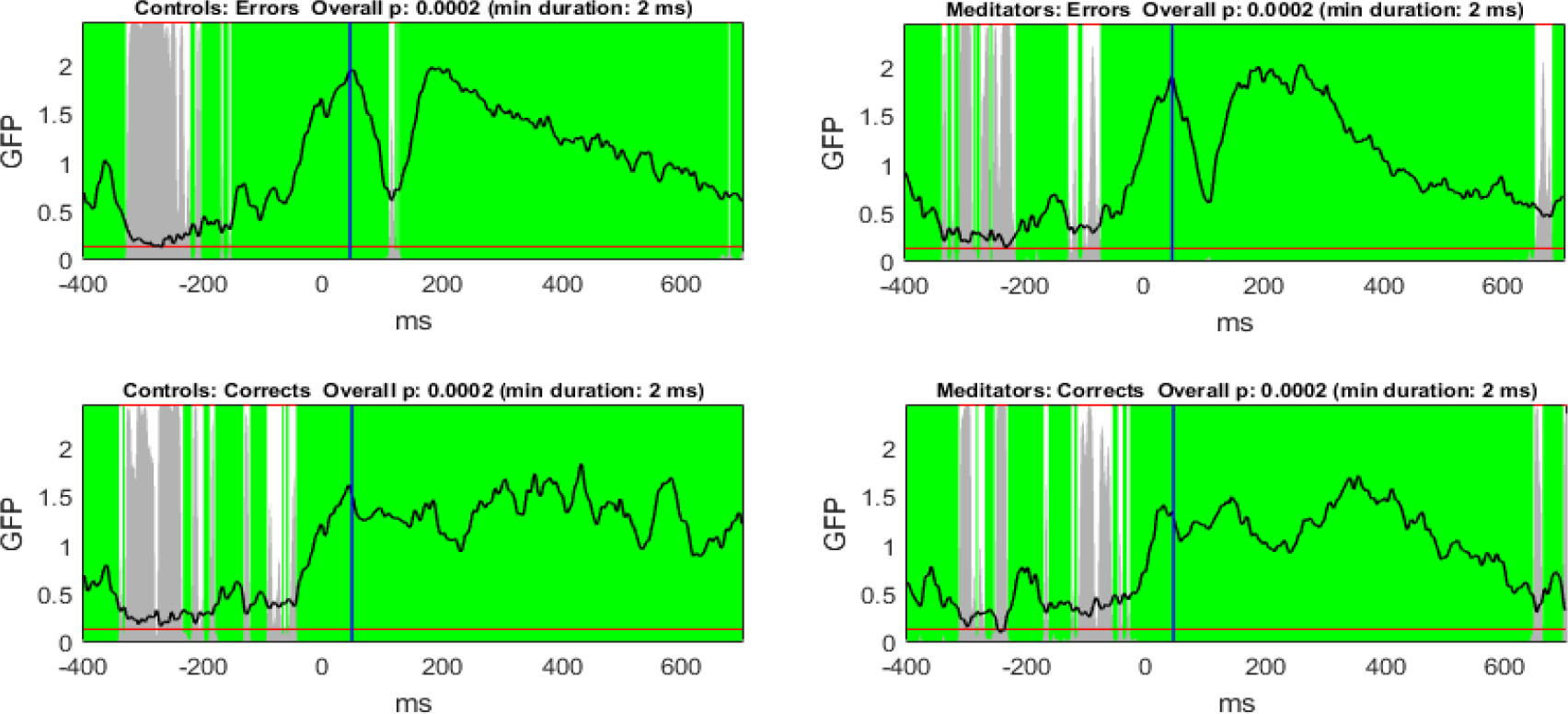
The TCT test showed consistent topographical activation across the epoch in each condition and each group. The black line reflects GFP, grey bars reflect the p-value (with the red line indicating p = 0.05), white periods reflect no significant differences between groups, and green periods reflect significant differences that survive duration multiple comparison controls.

### ERN and Pe TANOVA Showed No Differences Between Groups

No significant differences between groups were found in the TANOVA for the main effect of the comparison between groups nor for the group by correct/incorrect interaction (see figure 8). There was a significant difference for the main effect of trial type however, from immediately prior to the response until 700 ms post stimulus. These results indicate that that the distribution of neural activity did not differ between the meditator and control groups, and that no group differences were asymmetrically present for the different trial types at any point across the epoch (including during the ERN and Pe windows). The results do however indicate that error responses result in a different distribution of neural activity to correct responses from just prior to the response for the remainder of the epoch. Figure 9 shows the topography of activity averaged across both groups during the ERN and Pe periods for both error and correct trials. These topographies demonstrate the typical ERN and Pe patterns for error trials, but not for correct trials, indicating that the task did generate the expected neural activity during error trials.

**Figure 8.**
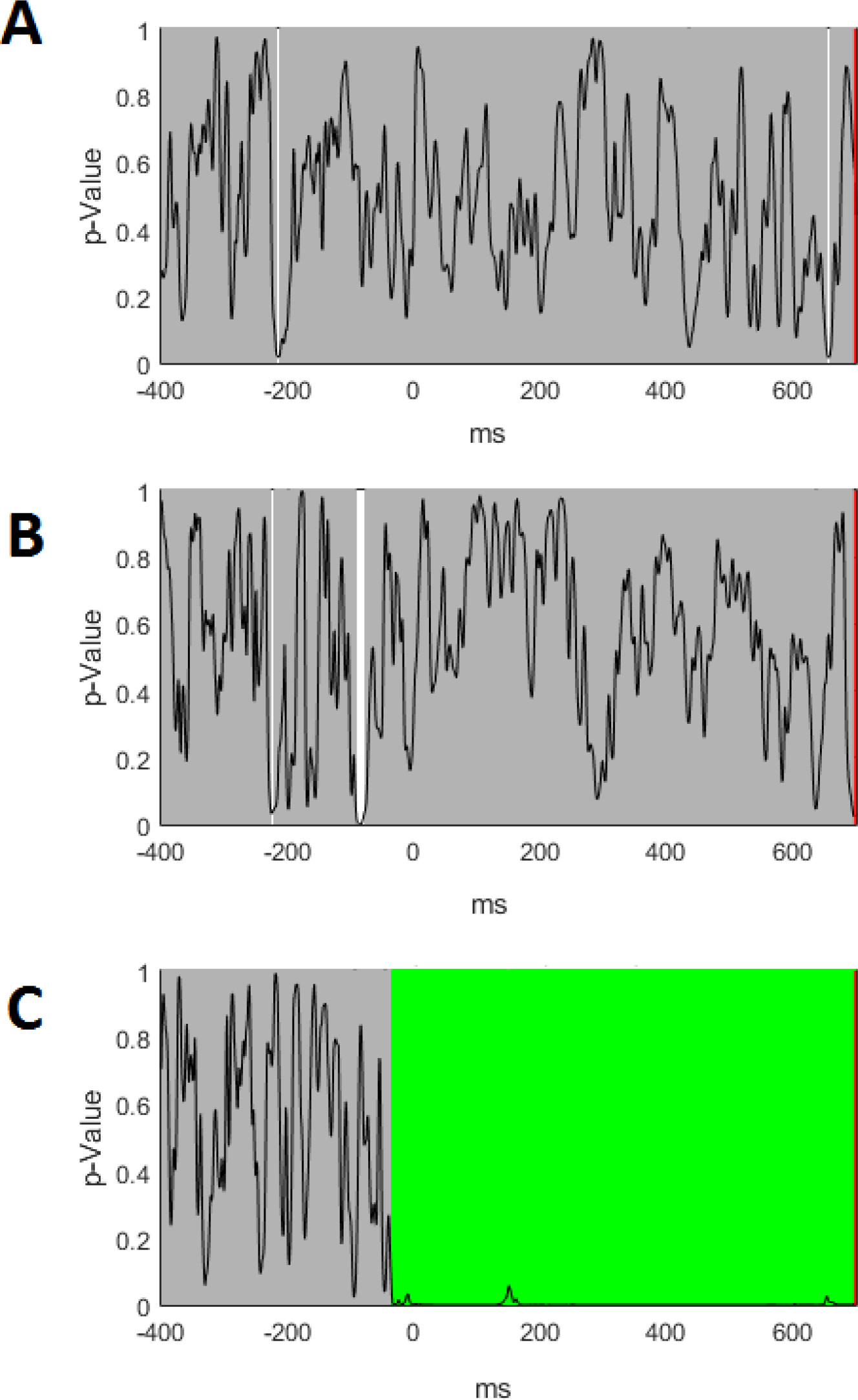
P-value graphs for TANOVA comparisons. The black line reflects the p-value, grey periods reflect no significant differences between groups, white periods reflect significant differences that did not survive duration multiple comparison controls, and green periods reflect significant differences that do survive duration multiple comparison controls. A - Group main effect, B - Group by correct/incorrect interaction, C - Correct/incorrect main effect (note the significant difference from −34 ms onwards, which survived duration control for multiple comparisons).

**Figure 9.**
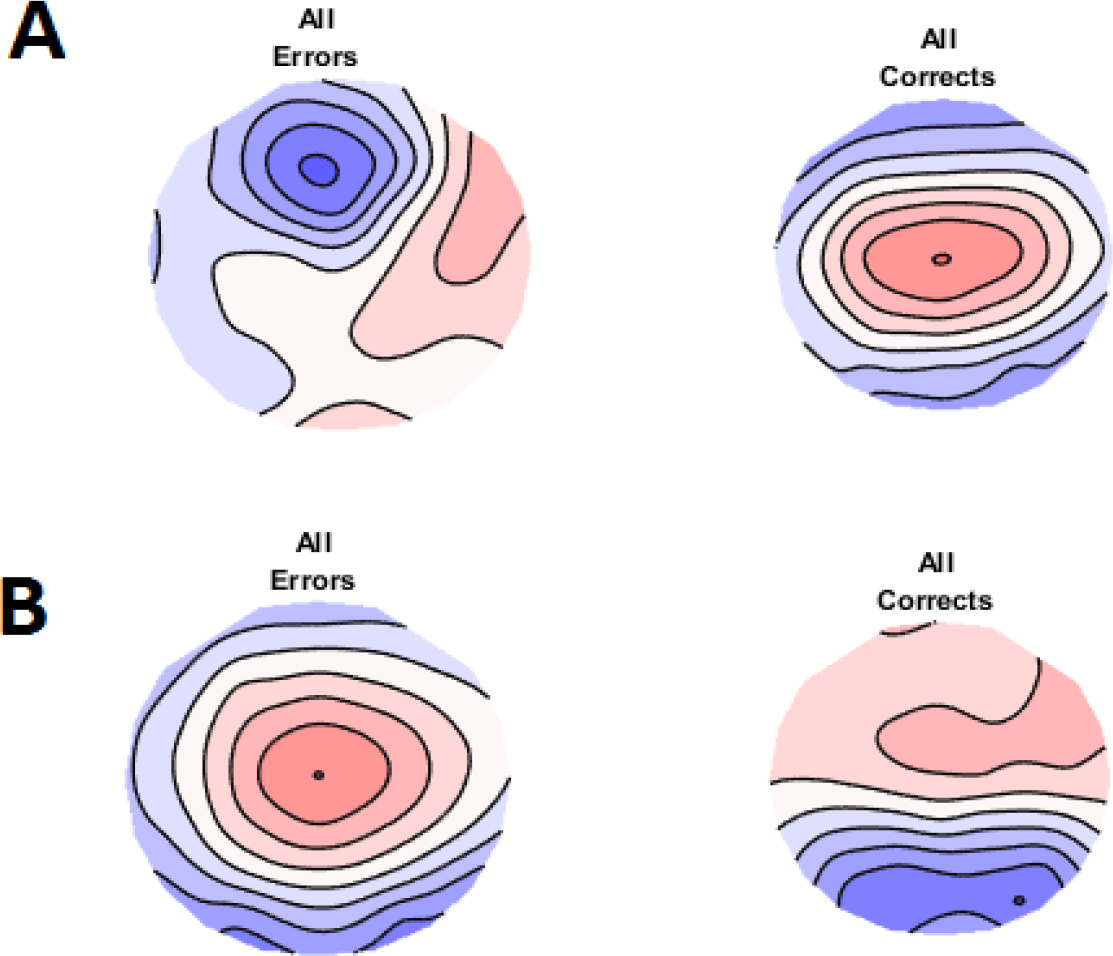
A - Averaged topography for the ERN 50 - 150 ms window across both groups for errors and correct trials. B - Averaged topography for the Pe 200 - 400 ms window across both groups for errors and correct trials.
Figure 10. Alpha activity GFP across both groups and across the epoch. A - GFP alpha comparisons between error and correct trials (the line reflects p-values, the green period reflects activity that significantly differs between conditions and survives duration multiple comparison controls). B - Averaged GFP alpha activity during the significant window (p < 0.001).

In order to ensure our results were not simply due to the specific parameters of measurements chosen, we also tested other potential baseline correction parameters (baseline correcting around the response from −200 to 0 ms, from −400 to −200 ms, and using the absolute baseline correction method to the entire epoch), and other criteria for the number of accepted epochs before participants were included (6 epochs required resulted in 29 meditators and 23 controls, 14 epochs required resulted in 12 participants in each group). We also computed difference ERPs by subtracting error related trials from correct related trials. Once these alternative EEG processing parameters had been computed, we performed the same statistical comparisons to those reported above (except for with the difference ERPs, which were compared using between group t-test designs in RAGU). None of these permutations altered the results currently reported (all p > 0.05).

### Alpha Suppression

Comparisons using RAGU indicated that error trials showed significantly less alpha power than correct trials from 155 ms until the end of the measured epoch when compared across both groups, indicating significant alpha suppression following error trials similar to Bing-Canar et al. (2016) (p < 0.001 when data was averaged across the significant window). Results of this comparison can be viewed in figure 10. However, although error trials showed the suppression effect across both groups, no significant differences or interactions were present between groups and correct / error trial alpha power GFP or TANOVAs (all p remained > 0.05 across the entire epoch). This result suggests that alpha suppression following error trials was not stronger in meditators compared to controls, in contrast to the results found by Bing-Canar et al. (2016).

**Figure 10.**
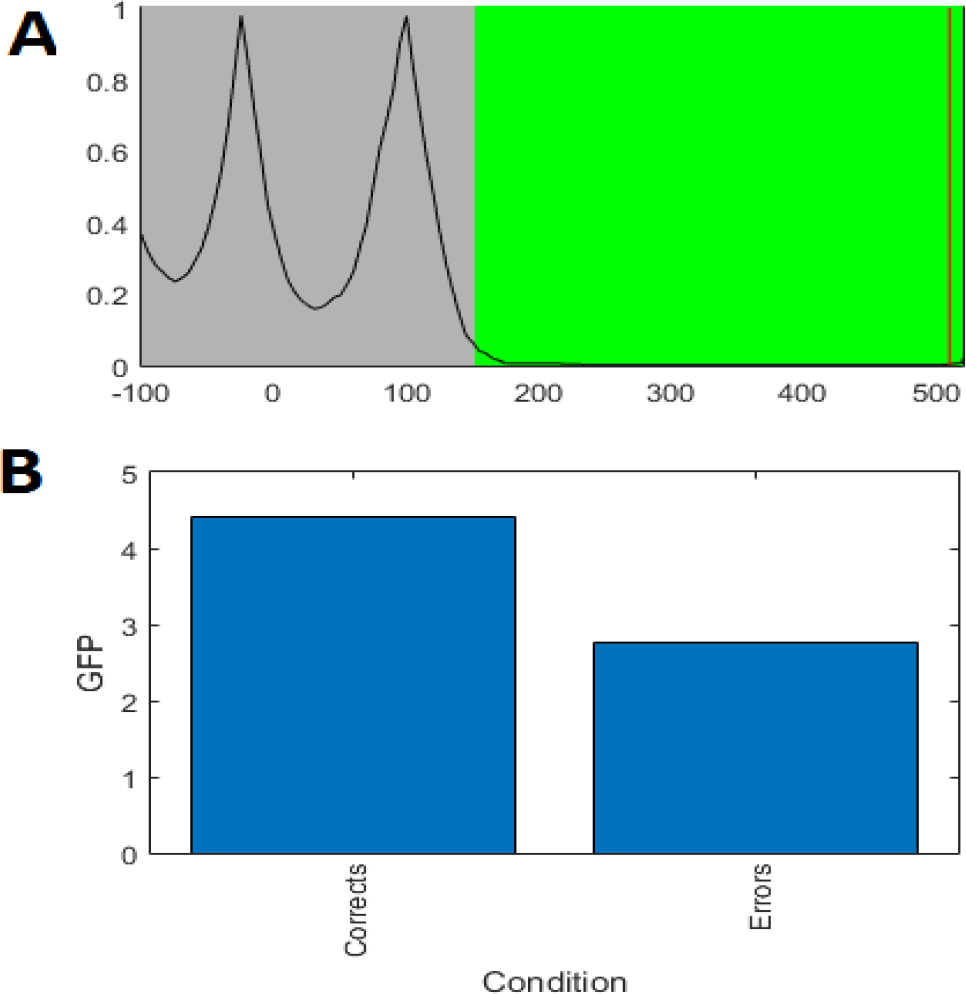
Alpha activity GFP across both groups and across the epoch. A - GFP alpha comparisons between error and correct trials (the line reflects p-values, the green period reflects activity that significantly differs
between conditions and survives duration multiple comparison controls). B - Averaged GFP alpha activity during the significant window (p < 0.001).

In order to ensure our results were not simply due to the specific parameters of measurement selected, we also compared alpha suppression between groups using Morlet Wavelet transforms, in order to determine if this would provide significance due to the method’s higher temporal resolution. Additionally, we computed difference activity by subtracting error trial alpha activity from correct trial alpha activity, and compared these values between groups in RAGU using a t-test design. Comparisons using the Morlet Wavelet transform or correct minus error trial alpha activity did not alter the results currently reported (all p > 0.05).

## Discussion

Our study showed evidence against differences between experienced mindfulness meditators and healthy control non-meditators in the ERN and Pe measures of error processing. Bayes Factor analysis suggested the strength of the evidence for the null hypotheses ranged from substantial to strong for these two ERPs (Jeffreys, 1961). Frequentist statistics also showed lack of evidence for differences in alpha suppression, although Bayes analyses of single electrode data showed weak support for stronger alpha power following correct responses in controls (of a size that is similar in strength to anecdotal evidence) (Jeffreys, 1961; Raftery, 1995). Analysis parameters for these measures vary between studies, so in order to confirm our null results were not study-specific due to the particulars of analysis parameters chosen, we analysed the data using measures that take into account all electrodes and timepoints (while controlling for multiple comparisons), as well as traditional single electrode analyses. We also performed analyses with multiple different baseline correction periods, and a range of accepted error epochs required for participant inclusion. None of these variations resulted in significant differences between groups. Post-hoc power analysis demonstrated our study was sufficiently powered to detect effect sizes similar to those found in previous research. This suggests our null result was not simply due to our particular analytic approach or a lack of statistical power.

### Potential explanations for the null results

There are a number of potential explanations for the current null results. One possible explanation is that our inclusion criteria of mindfulness meditators was too broad, resulting in a diversity of practice types that obscured differences associated with any specific practice. Notably, however, our recruitment methods and inclusion/exclusion criteria were quite comparable to those used by Teper and Inzlicht (2013). Thus, while we cannot out this lack of specificity, similar methods yielded significant results in the past. Additionally, the TCT test demonstrated consistent neural activity within the meditation group, suggesting common neural activity within the group rather than potential diversity of practice leading to diversity of neural activity. Another possibility is that the null results are specific to the group of meditators sampled, and previous studies accurately conclude that meditators differ in neural activity related to error processing. However, meditators in the current study self-reported higher levels of mindfulness on both the FMI and FFMQ, and showed differences in reacting to errors in a different fashion in the two FMI questions that could be viewed as most relating to subjective error processing. This suggests that meditators subjectively experience processing errors in a different manner. Neural activity is commonly accepted to be related to subjective experience, so we would expect these differences to be reflected in neural activity. Meditators in the current study did show differences in Stroop or Go/Nogo stimulus locked neural activity, indicating differences neural activity were present, just not error processing related neural activity (Bailey et al in preparation, Raj et al in preparation). Of note, our meditation group had the greatest experience in terms of number of years practicing meditation of any study of error processing conducted so far (mean = 8.72, compared to Andreu et al. 2017, mean = 5.1). As such, we expected that if general mindfulness meditation practice were to affect EEG measures of error processing, the current study should have shown stronger differences than previous studies.

Given the differences in self-reported experience of error processing, there are two possible explanations that seem more likely than the above. Firstly, that meditators may differ in neural processing of errors, but that EEG is an insufficient measurement tool to capture these differences. Our results indicated that meditators showed the same neural activity as controls in terms of the neural regions engaged (the distribution of neural activity did not differ between groups) and the strength of activity (neural response strength reflected by GFP did not differ between groups). However, each EEG electrode measures the summed post-synaptic potentials of millions of neurons from a cubic centimetre of cortex, and only the neurons oriented towards the scalp and on the outer layer of the gyri (Buzsáki, Anastassiou, & Koch, 2012). As such, EEG measures could not detect differences in activation in deeper sources, or differences in the particular neurons activated if those neurons were still located within the same cubic centimetre of cortex. fMRI research has shown differences in meditators in the activation of deeper sources such as the ACC, ventrolateral prefrontal cortex and the insula in association with response inhibition and negative valence processing, so this may be a potential avenue to examine whether meditation effects error processing in deeper brain (Allen et al., 2012). Additionally, it may be that differences in error processing in meditators are present, but only at a longer delay following the error than the 700 ms measured in the current study. Perhaps the attentional and emotional regulation mechanisms engaged in the meditation group occur over the time-scale of seconds, in which case the current EEG measures would miss any potential differences. This proposed slower difference in error processing aligns with the lack of difference in post-error slowing (found in both the current research and by Andreu et al. 2017). The differential processing of errors in meditators may occur too slowly to adjust behavioural performance in the next trial. This explanation accounts for the self-reported subjective differences in reactions to errors, as well as the current null results. However, future research would need to provide evidence for differences measured with tools other than EEG, or at longer time scales than those measured in the current study before this explanation would be viable, particularly in the absence of objective behavioural differences such as post-error slowing.

The second potential explanation is that there is no difference between meditators and controls in error processing, in contrast to the conclusions of previous research. This explanation conflicts with self-reports of meditators which indicate that they experience error processing in a different fashion to non-meditators. It may be that the self-report data from meditators is not an accurate reflection of their error processing. In that case, this explanation is the most likely given the current data. Unfortunately, the current results provide no way to discriminate between overall lack of difference, and lack of ability to detect differences via an objective method. Arguably, though, the lack of an objective indicator suggests a lack of a difference, pending further evidence that could support subjective perceptions of a difference in error processing in meditators.

### Comparison with previous research

If the explanations suggesting there are no differences in EEG measures of error processing are correct, the positive results of previous research require explanation. We suggest the positive results of previous research could be due to a number of factors. It may be that meditation resolves abnormalities in error processing in clinical populations, for example by increasing previously blunted ERN amplitudes (Smart and Segalowitz 2016). However, the results of the two studies in clinical populations conflict. Fissler et al (2017) indicated an increase in blunted ERN amplitudes in depression, while Schoenberg et al (2014) found ADHD participants showed a reduction in ERN amplitude (even though ERN amplitudes are already reduced in ADHD - Geburek et al 2013). There may also be a non-linear relationship between meditation experience and EEG measures of error processing. However, if this is the case it is not clear why differences would be apparent after 15 minutes (as per Bing-Canar 2016, Larson 2013, and Saunders 2016) but then return to baseline after the average of eight years of experience in the current study. Further research using more rigorous methodology in clinical populations and a range of meditation experience will be required to resolve these points.

Another potential explanation for the conflict between the current null results and the positive results of previous research is that a wide range of analysis parameters are available for selection, and the choice of specific parameters for analysis can positively bias the results (Kilner 2013 - see supplementary methods for a discussion of these parameters). Lastly, it is possible that the inconsistent results of previous research and the current null results are due to chance variation. Chance variation seems a likely explanation, because previous results have been inconsistent (see Table 1 and summary in supplementary materials). The explanation that accounts for the largest amount of the literature (including the current null result) is a combination of the choices of specific analysis parameters, chance variations between studies, and potentially that meditation resolves pathological error processing in clinical populations (although this requires further research with more rigorous methodology to confirm). It is also possible that any real effects of meditation practice are smaller than previously suggested, which would align with other prominent replication failures in the psychological literature.

The two previous studies to examine error processing in long term meditators both showed higher ERN amplitudes (Teper and Inzlicht, 2012, Andreu et al. 2017). Andreu et al. (2017) found an enhanced ERN and CRN, suggesting the difference is not specific to error processing but occurs following correct responses also (while Teper and Inzlicht 2012 did not include correct trials in their analysis). Andreu et al.’s (2017) research was specific to Vipassana meditation, while our sample included a broad spectrum of practices (which all fit under the Kabat-Zinn (1994) definition of “mindfulness meditation”). Further research is required to determine whether the current null results, the previous positive results, or something in between most accurately reflects neural activity related to error processing in experienced meditators, and whether differences may be specific to particular practices.

### Interpretation

What do these results mean for decisions regarding whether to practice mindfulness meditation, or whether mindfulness is clinically applicable? Firstly, mindfulness meditation is not a panacea - while it shows positive effects on some processes (Kerr et al., 2011; Lutz et al., 2009; Slagter et al., 2007), it seems unlikely that it will have an impact on all measures (as per the lack of an effect on error processing in the current study). As such, if the findings in the current study are replicated, the results could suggest that mindfulness meditation may not be a helpful treatment for illnesses where the neurophysiology underlying error processing seems to be a primary impairment (if future research were to demonstrate that to be the case for a particular illness). However, more work is required to determine whether mindfulness meditation resolves error processing abnormalities in clinical populations. In particular, the ERN has been suggested to be a potential biomarker for anxiety (Meyer, 2016), for which meta-analysis has suggested mindfulness-based interventions are an effective treatment (Vøllestad, Nielsen, & Nielsen, 2012). Additionally, even if meditation does not alter error processing, previous research has suggested it is likely to have an effect on other neurophysiological measures such as the P3. Moreover, these differences have been shown to relate to improved behavioural performance (including in the current sample of participants - Bailey et al. in submission). The P3 has been indicated to be abnormal in a range of psychiatric illnesses, so despite the null results in the current study, meditation may still be helpful in resolving abnormalities in neural activity (Bostanov, Keune, Kotchoubey, & Hautzinger, 2012; Schoenberg et al., 2014). Ultimately, considerably more work is needed to identify which neural processes mindfulness meditation impacts and the clinical implications of those processes (cf. Van Dam et al. 2018).

Our results also have implications for the understanding of the potential mechanisms of meditation. If future research shows that our results do accurately indicate the lack of an effect of meditation on EEG measures of error processing, the results might allow for inferences regarding brain regions and neurocognitive processes that are not affected by meditation. In particular, the ERN is thought to be generated by the theta activity in the ACC (Dehaene et al., 1994). A comprehensive framework of the ERN suggests it reflects an automatic tagging of phasic ACC activity with a negative or positive valence, which reflects alignment or lack of alignment of behaviour with goals (Dehaene et al., 1994). The current results suggest this automatic process may be unaltered by meditation. In alignment with this interpretation, the meditators in the current sample also showed no differences in the Nogo N2 to stimulus locked activity (Bailey et al. in preparation) (an ERP also generated by the ACC and thought to reflect theta modulation (Cavanagh, Zambrano-Vazquez, & Allen, 2012)). However, it should be noted that other research has shown differences between meditators and controls in the N2 and in theta activity (Aftanas & Golosheykin, 2005; Cheng, Chang, Han, & Lee, 2017; Sanger & Dorjee, 2016; Xue, Tang, Tang, & Posner, 2014). Our suggested explanation for the variation in result between studies is that meditation may increase the ability to dynamically modulate neural oscillations (with both increases and potentially decreases across multiple frequencies), when frequency modulations are beneficial to meeting task demands or goals, rather than ubiquitous increases in specific oscillation frequencies (Bailey et al. in preparation).

Similarly, the Pe is thought to be generated by the cingulate cortex and the insula (Herrmann et al., 2004; O’connell et al., 2007; Overbeek et al., 2005; Ullsperger et al., 2010; Vocat et al., 2008). These regions have been shown to be altered in meditators (Fox et al., 2014). The Pe is also thought to reflect awareness of the error or attention towards the motivational factors associated with the error (Endrass et al., 2012; Hughes & Yeung, 2011; Nieuwenhuis et al., 2001; Ridderinkhof et al., 2009; Shalgi et al., 2009). These factors seem likely be altered as a result of meditation, which is thought to affect both attention and awareness (Jha et al., 2007; Tang et al., 2007; Teper et al., 2013). Additionally, the Pe has been suggested to reflect a P3b for affective processing related to an error (Davies et al., 2001; Overbeek et al., 2005). Both emotional regulation and the P3b seem to be affected by meditation, including in the current sample (Aftanas and Golosheykin 2005, Schoenberg, Hepark et al. 2014, Bailey et al. in preparation, Raj et al. in preparation). As such, a comprehensive interpretation of the current results and previous literature would suggest that although the P3b is altered by meditation (along with related cognitive processes and brain regions), the alteration does not extend to the processing of errors.

Lastly, the frequentist results indicated no evidence for differences in alpha suppression following errors in the meditation group, while the Bayesian results of analyses including only selected electrodes indicated weak support for differences in alpha suppression (of a similar strength to anecdotal evidence) (Jeffreys, 1961; Raftery, 1995). Post-error alpha suppression is suggested to reduce neural idling and increase vigilance (Carp & Compton, 2009). Enhanced modulation of alpha activity in meditators has been shown in previous research (Kerr et al., 2011). Since meditation is thought to increase the ability to attend, it might be expected that meditators would show an enhanced ability to modulate alpha following an error. However, it has been suggested that post-error alpha suppression is a non-adaptive response to errors, reflecting increased vigilance but not increased behavioural control, as it has not been associated with improved performance (Schroder & Infantolino, 2013). In the absence of further evidence, the reasons for the current null (or at most, very small effect size) results are unclear. It may be that meditators do not show differences in error processing that occur early enough for post-alpha suppression to be modulated, which may similarly explain the lack of altered post-error reaction time slowing in the meditation group. However, the data depicted by figure 10 suggests that while controls show alpha suppression following errors, meditators show low alpha following both correct and erroneous responses. This may suggest more constant vigilance, rather than increased vigilance only following errors. This interpretation should be viewed with caution however, as it was only weakly supported with Bayesian statistics, which only included selected electrodes.

### Limitations, and Future Research

Although the current study included a well-matched control group (in terms of gender and age) the meditation group showed a higher level of education. However, the effect of this confound is to increase the chance that the current study would obtain a positive result (although not due to the intended factor). As such, the difference in education is not likely to be a limiting factor in interpretation of the current results. Similarly, the lack of a cross-sectional design would have been a limiting factor in drawing conclusions regarding causation. However, in context of null results, the significant experience of the meditators recruited is a strength of the study that would have been very difficult to achieve with a longitudinal design. Another strength of the current research is that it showed neural activity was consistent within groups and that the typical patterns of neural activity were obtained in all measures (including differences in ERN topography between correct and incorrect trials, enhanced Pe amplitude and a Pe specific topography in error trials, and enhanced alpha suppression following error trials). As such we are confident that the tasks and EEG processing steps used generated valid error processing related activity for comparison between groups.

Additionally, although controls showed a longer RT for correct compared to incorrect responses to the emotional Stroop, correct trials were only included in the analyses to determine whether the difference in neural activity between correct and error trials was larger in one of the groups compared to the other. The error related neural activities were the main variable of interest, and since neither group differences nor interactions between group and trial type were significant in the neural analyses, we suggest that the difference in correct trial reaction time in the emotional Stroop task would not limit the conclusions drawn from the lack of between group differences.

Another potential limitation is that the parameters for measuring error processing have not been well defined by previous research. In particular, there is no consensus on the number of accepted error trials before a participant should be included in analysis. We have taken a conservative approach and used a minimum of 10 accepted error trials, which is the number at which the standard deviation between trials stabilizes and high internal reliability measured with Cronbach’s alpha is obtained (Olvet & Hajcak, 2009). More commonly, research examining error processing has used 5-8 error trials, which some researchers have suggested is too few (Fischer et al., 2017; Larson et al., 2010). However, the current null results were unchanged even after re-analysis including participants with only 6 accepted error trials.

Lastly, although the sample size of the current study could be considered moderate (22 meditators, 20 controls) our sample had sufficient power (> 0.85) to detect effect sizes similar to those shown by previous research (Teper and Inzlicht, 2012). Additionally, even when only 6 error epochs were required for analysis (providing 29 meditators and 23 controls) our null results were unchanged. Bayes Factor analysis was also used, in order to provide substantial evidence in support of the null hypothesis for the ERN and Pe (Jeffreys 1961). All these lines of evidence suggest evidence for a null result, which is unlikely to be simply due to insufficient power, and more likely reflects a genuine finding.

Overall, the current results add a necessary discernment to the research surrounding mindfulness meditation. Given the current hype surrounding the practice (Van Dam et al 2018), it is important to delineate not only the positive effects of the practice, but the areas which are unaffected. While mindfulness may have positive behavioural and physiological consequences, it cannot be assumed to improve all components of attention and self-regulation. This increased discernment will enhance our understanding of the effects and uses of mindfulness practice in the future.

### Data Availability Statement

Participants involved in the study did not provide consent to data sharing, and data sharing was not approved by the ethics committee, so the results reported in the paper comprise the complete data available for sharing

### Compliance with Ethical Standards

All procedures performed in the study involving human participants were in accordance with the ethical standards of the institutional research committee and with the 1964 Helsinki declaration and its later amendments. Informed consent was obtained from all individual participants included in the study.

## Supplementary Materials

### Summary of the inconsistencies of previous research

With regards to the ERN, two studies have shown no differences in university students after a single 15 minute mindfulness training session (Larson et al. 2013, Bing-Canar 2016). Four studies have reported increases in ERN, or in the difference between error and correct ERN amplitude (one in in university students after a single 15 minute mindfulness training session, one in depression after a two week intervention, one in healthy aging adults after eight weeks, and one in long-term meditators with a cross sectional design) (Saunders 2016, Fissler 2017, Smart 2017, Teper 2012). One study also reported a decrease in the difference between error and correct ERN amplitude (in ADHD participants after a twelve week intervention) (Schoenberg et al 2014). Results for the Pe are even more suggestive that the overall result is that meditation does not have an effect on this process. Only two studies indicate that meditation had an effect on the Pe. One study showed that after 15 minutes of mindful breathing, university students displayed smaller Pe amplitudes than the control group (Larson et al 2013), and the other showed that 12 weeks of mindfulness increased the Pe in ADHD (Schoenberg et al 2014). All other studies showed null results, similar to the current research. Lastly, our results also indicated a lack of difference in alpha suppression following errors in the meditation group, in contrast to Bing-Canar et al (2016). However, Bing-Canar’s (2016) results are likely to reflect a state effect after only a single 15 minute session of mindfulness practice, so in the absence of further research on alpha suppression in meditation, we would conclude that there is no trait effect on alpha suppression from mindfulness meditation.

We recommend that future research prevent potential experimenter influence on results by using analysis methods that include all available electrodes and timepoints. Additionally, we recommend that in the absence of validated baseline correction parameters or consensus on the number of required epochs for valid inclusion in analysis, multiple options for each of these parameters be tested and reported. This will ensure the specific parameters selected do not bias the results. Lastly, we recommend that if difference curves between error and correct neural activity are calculated, that separate analyses including both trial types in repeated measures designs are also reported (along with means and standard deviations), to ensure results are not specific only to the interaction between group and correct / incorrect trial response, or if the results are specific to the interaction, that information is made clear to future researchers. These methods will also allow for more effective meta-analyses to be performed.

### Additional minor potential limitations

The use of errors from different tasks may have introduced an unforeseen confound. In particular, reaction time differences were present between error and correct trials in the control group but not the meditation group. This may have introduced a between group confound that was unaccounted for by the study design. However, the different reaction times in controls would have had the effect of introducing a different pattern of background stimulus related neural processing between response locked error and correct trials. Group differences in this effect would have enhanced the potential differences between groups, so the difference in reaction times between correct and incorrect trials in the control group is not a confound in the context of a null result. Additionally, the neural activity from the stimulus that would have been introduced in this fashion is likely to be variable due to the variability of reaction times. As such, any effect is likely to have averaged to zero. Previous research has also indicated error related activity is similar across different tasks, so combining tasks is unlikely to be an explanation for the null result (Riesel, Weinberg, Endrass, Meyer, & Hajcak, 2013). This is supported by the consistent pattern of neural activity within groups shown by the TCT test in the current study.

A large number of participants were excluded due to an insufficient number of accepted error epochs. This is fairly typical of error related processing research (Fischer et al., 2017; Larson et al., 2010; Pontifex et al., 2010). However, the exclusion does elicit concerns about generalisability. Error processing potentials correlate with number of errors (Fischer et al., 2017). As such, excluding high performing participants from both groups means the current study did not sample the full distribution of error processing related neural activity. Future research should design tasks of sufficient difficulty and numbers of trials so that all participants produce enough errors to be included in the final analysis.

Author Contributions
NWB designed and oversaw the study, assisted with data collection, performed the data analysis, and wrote the paper. KR performed data collection, assisted with data analysis, and collaborated with writing of the study. GF performed data collection, assisted with data analysis, and collaborated with writing of the study. BMF assisted with study design and writing of the paper. NCR assisted with study design and writing of the paper. NTVD collaborated in the interpretation of results, writing and editing of the final manuscript. PBF assisted with study design and writing and editing of the final manuscript.

## Abbreviations

ERN: Error Related Negativity
Pe: Error Positivity
EEG: Electroencephalography
MBSR: Mindfulness Based Stress Reduction
MBCT: Mindfulness Based Cognitive Therapy
FMI: Freiburg Mindfulness Inventory
FFMQ: Five Facet Mindfulness Questionnaire
ACC: Anterior Cingulate Corte
DLPFC: Dorsolateral Prefrontal Cortex
ADHD: Attention-Deficit Hyperactivity Disorder
BAI: Beck Anxiety Inventory
BDI-II: Beck Depression Inventory II
ICA: Independent Component Analysis
AMICA: Adaptive Mixture Independent Component Analysis
RAGU: Randomisation Graphical User Interface
TCT: Topographical Consistency Test
TANOVA: Topographical Analysis of Variance
GFP: Global Field Potential
RMS: Root Mean Squared
FFT: Fast Fourier Transform

